# A versatile microfluidic extrusion-based hydrogel platform for self-organization and long-term maintenance of engineered 3D lymphatic endothelium

**DOI:** 10.1101/2025.08.21.666813

**Authors:** E. Mazari-Arrighi, A. Boyreau, L. Chaillot, W. Souleyreau, L. Andrique, T. Mathivet, A. Bikfalvi, P. Nassoy

## Abstract

Lymphatic endothelium is essential for interstitial fluid drainage, immune surveillance, and macromolecular transport. Existing *in vitro* models that accurately recapitulate its three-dimensional structure, barrier properties, and long-term stability are still limited. In this study, we propose a robust, adaptable, and scalable platform designed to generate engineered three-dimensional lymphatic endothelium using primary human dermal lymphatic endothelial cells (HDLECs). This system combines microfluidic extrusion-based fabrication allowing precise control of the tubular geometry with custom-optimized extracellular matrix–derived hydrogel. The capability to tune dimensions enables the creation of constructs that encompass physiologically relevant size ranges. Through a systematic matrix screen, we identified a unique four-component formulation—gelatin, Matrigel, hyaluronic acid, and fibrinogen—that supports the rapid self-assembly of HDLECs into stable, lumen-forming monolayers within one week. These engineered structures maintain viability and structural integrity for a minimum of 30 days under static culture conditions. Functional permeability assays demonstrated selective tracer uptake characteristics of lymphatic transport. Comparative studies with blood vascular endothelial cells indicate that the proposed platform preserves maintenance of lineage-specific expression profiles, junctional organization, and permeability properties under identical fabrication parameters. Altogether, this approach offers a reproducible and controlled system for studying three-dimensional endothelial architecture, transport mechanisms, and extracellular matrix remodeling across diverse endothelial phenotypes, thereby addressing a pivotal gap in the modeling of lymphatic vasculature.

**Highlights:**

- A microfluidic extrusion platform allows primary human lymphatic endothelial cells to self-organize into 3D hollow tubes.
- The internal dimensions of the hybrid cell/hydrogel constructs can be precisely controlled by tuning the flow rates.
- A custom-defined four-component extracellular matrix provides essential biochemical cues for stable monolayer formation and for the in vitro organization of lymphatic endothelial vessels.
- Matrix composition significantly influences cell proliferation and migration, thereby impacting on the spatial arrangement of the tissue.
- The three-dimensional lymphatic tissue configuration sustains cell viability, consistent expression of lineage-specific markers, and barrier function which lasts for at least 30 days.
- The system maintains morphological and functional properties not only for lymphatic endothelial cells, but also for blood endothelial cells in long-term culture conditions.

## Introduction

The vascular system is composed of two complementary networks—blood and lymphatic vessels—that coordinate to regulate fluid balance, orchestrate immune surveillance, and sustain tissue homeostasis [1]. Importantly, these two networks are built upon a shared anatomical blueprint: highly polarized endothelial monolayers encasing a central lumen, organized into hierarchically branched architectures that span from micrometer-scale capillaries to millimeter-scale conduits [1]. Together, they form a contiguous three-dimensional endothelial interface distributed throughout the body [2, 3].

Lymphatic endothelial cells (LECs) originate from a specific developmental lineage and are regulated by distinct transcriptional, metabolic, and signaling mechanisms[4– 6]. Compared to their blood vascular counterparts, LECs form vessels characterized by greater permeability and are uniquely equipped for macromolecular uptake and interstitial fluid drainage. This transport capacity is central to lymphatic physiology and underlies the roles of the lymphatic vasculature in immune cell trafficking, inflammation, fibrosis, and tumor progression [6–9] .

The development of the lymphatic vasculature during embryogenesis has been extensively investigated using *in vivo* models, particularly in mice and zebrafish [10, 11]. Nevertheless, reproducing this context-dependent behavior *in vitro* remains a major experimental challenge ([4, 12]). Animal models are constrained by interspecies differences and limited scalability. Two-dimensional cultures lack physiological architecture and provide limited insight into spatial organization and mechanical cues relevant to lymphatic endothelium [12]. Three-dimensional systems have recently addressed key physiological features of lymphatic endothelium [13], such as permeability [14–16], intercellular junctions [17–19], or sprouting behavior [20–22], with some incorporating flow or matrix barriers to mimic interstitial drainage [23]. However, many of these models are short-lived, typically limited to less than one week of culture, thereby limiting the potentiality to explore how the lymphatic endothelium organizes and reaches homeostasis within a physiological tissue environment. Additionally, several engineered systems rely on lymphatic-like cells derived from pluripotent stem cells, which often lack the maturity and stability of primary adult LECs [24–26].

In this work, we implemented a microfluidic extrusion-based strategy to produce hydrogel tubes seeded with primary human LECs, in which the microenvironmental composition can be adequately controlled. This approach offers a scalable and reproducible method to generate large numbers of highly uniform hydrogel tubes, enabling systematic studies with high experimental throughput. These constructs replicate key features of 3D lymphatic endothelial vessels by precisely controlling their size and the tubular organization. By adjusting construct size and matrix composition, we determined the exact conditions for lymphatic cells to self-organize in 3-dimensions. These conditions support remodeling of the initial hydrogel matrix by the seeded cells, maintain the expression of key lymphatic endothelial markers, and preserve selective permeability for at least 30 days of culture. The platform is also compatible with blood endothelial cells, enabling direct comparison of lymphatic and blood endothelial behavior within identical, size-controlled microenvironments. Altogether, these advances provide a platform uniquely suited to study lymphatic endothelial biology in defined and tunable microenvironments.

## Results

### 1. Coaxial extrusion produces size-controlled hydrogel tubes that support lymphatic endothelial self-organization

We used a coaxial extrusion method, adapted from capsule-forming hydrogel techniques [27], to create 3D lymphatic endothelium *in vitro*. This approach, previously applied in mechanobiology for cell confinement [28–30], enables precise control of the morphogenic process. Primary human dermal lymphatic endothelial cells (HDLECs) were suspended at 10^6^ cells/mL in a defined extracellular matrix composed of gelatin (2% w/v); Matrigel (30% w/v); hyaluronic acid (0.2% w/v); and fibrinogen (0.2% w/v). This mixture served as a core suspension that was co-extruded with a sorbitol-based intermediate solution and an outer sodium alginate solution. By entering into a calcium chloride bath, the alginate shell gelified to form stable hydrogel tubes (Fig.□1A).

**Figure 1.**
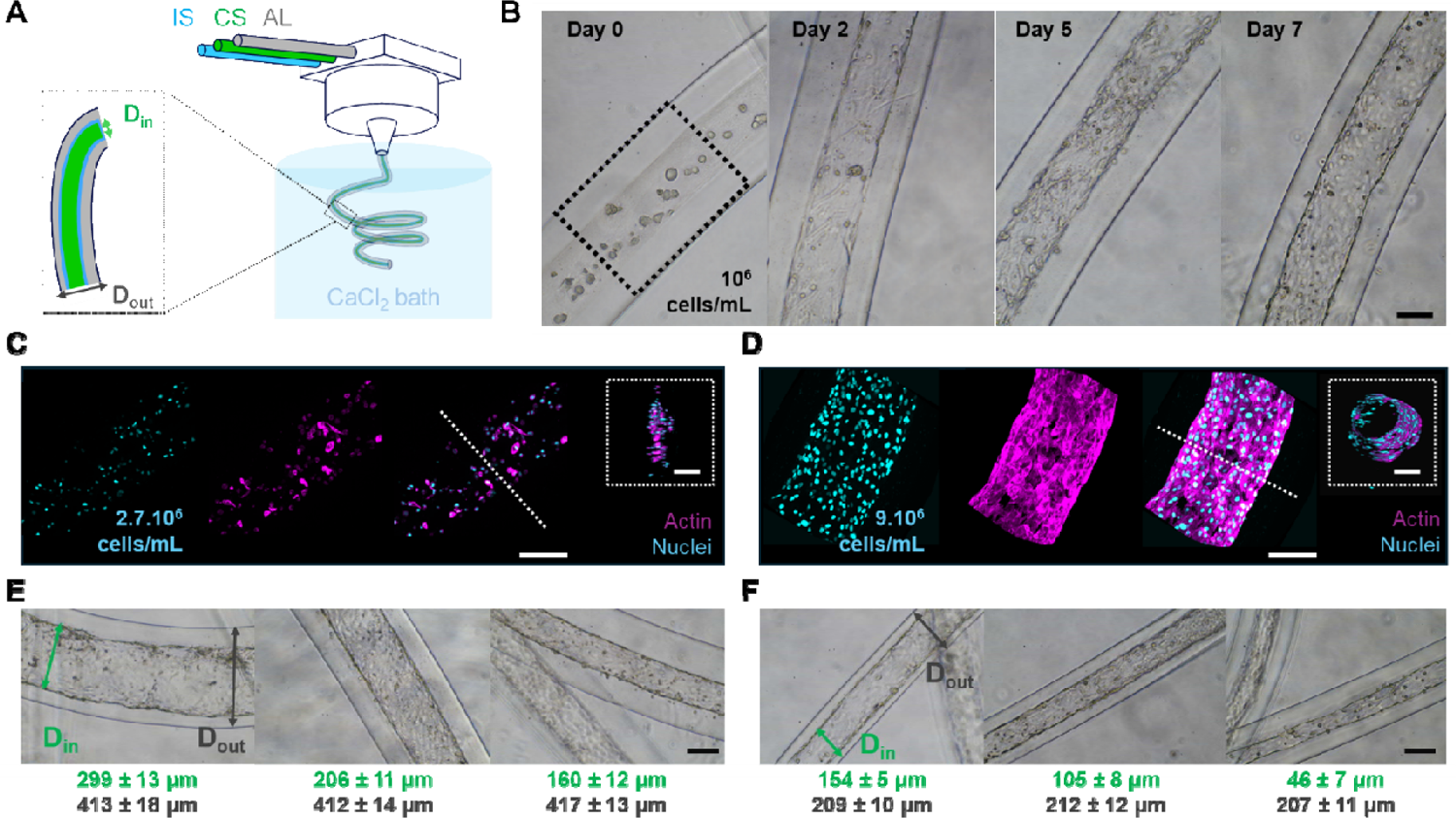
Tunable fabrication and early formation of in vitro 3D lymphatic endothelium. **(A)** Schematic of the coaxial micro-extrusion process. Human dermal lymphatic endothelial cells (HDLECs), suspended at 10□ cells/mL in a matrix composed of gelatin (G, 2% w/v); Matrigel (MG, 30% w/v); hyaluronic acid (HA, 0.2% w/v); and fibrinogen (FG, 0.2% w/v), are loaded into the core stream (CS). The CS is co-extruded with a sorbitol-based intermediate stream (IS) and an alginate shell stream (AL) into a calcium chloride (CaCl_2_) bath to crosslink the alginate and form stable, hollow hydrogel tubes. **(B)** Brightfield images at days 0, 2, 5, and 7 show the progressive redistribution, proliferation, and migration of HDLECs within the core compartments, which have an internal diameter of 200 µm. At day 0, a cell density of 10□ cells/mL corresponds to approximately 15 cells counted within a 500 µm-long segment outlined by the dashed light blue rectangle. **(C–D)** Confocal fluorescence images of constructs at day 2 **(C)** and day 7 **(D)**, showing actin (magenta) and nuclei (cyan). At day 2, cells remain sparse and mostly localized within the central hydrogel core. By day 7, they have proliferated and migrated toward the hydrogel–alginate interface, forming a continuous monolayer in a 3D configuration that conformed to the surface of the alginate tube. Insets show 3D reconstructions with selected viewing angles that highlight this transition from a central cell distribution to peripheral colonization along the alginate wall, resulting in a lumenized structure with no remaining cells in the core. **(E–F)** Brightfield images illustrating internal (D_in_) and external (D_out_) diameters achieved with distinct nozzle geometries. The core stream (CS) contained the cell-laden hydrogel, the intermediate stream (IS) consisted of the sorbitol solution, and the alginate stream (AL) formed the outer hydrogel layer. **(E)** With a nozzle of 450 µm inner diameter, tubes were extruded using CS/IS/AL flow rate combinations of 0.25/0.25/3.5, 0.5/0.5/3, and 1/1/2 mL/h (from left to right). **(F)** With a nozzle of 200 µm inner diameter, the corresponding combinations were 0.15/0.15/3.7, 0.5/0.5/3, and 1/1/2 mL/h (from left to right). D_in_ and D_out_ were quantified at three positions per tube (≥ 1 mm apart) across ≥ 3 independently extruded tubes per condition. Values are presented as mean ± standard deviation (SD). **Scale bars:** 100 µm.

Over a 7-day culture period, brightfield imaging revealed a progressive distribution of cells within the 200 µm-diameter core compartment. At day 0, cell density was approximately 10□ cells/mL (about 15 cells within a 200 µm diameter, 500 µm long cylinder), rising to a dense accumulation along the alginate wall by day 7 (Fig. 1B). Confocal microscopy within the hydrogel core showed that cell density increased to 2.7 × 10□ cells/mL (i.e. 40 cells counted in the same volume) at day 2, with cells still dispersed throughout the hydrogel core (Fig. 1C). By day 7, density reached 9 × 10□ cells/mL (i.e. 135 cells in the same volume), with cells forming a continuous monolayer along the alginate wall, and exhibiting fairly homogeneous distribution of cell nuclei and cortical actin enrichment (Fig. 1D). These observations were reinforced by 3D reconstructions from confocal stacks, which demonstrated complete cellular coverage of the lumenized structure. Note that the observed increase in cell density is consistent with a proliferation rate corresponding to a doubling time of 30– 40 hours [31–33].

Control of tube diameter was simply achieved by varying the flow rates set by the syringe pumps. In practice, the internal diameters were tuned by adjusting the relative flow rates of the core (CS), intermediate (IS), and alginate (AL) streams, while maintaining a constant total flow of 4 mL/h. With a 450 µm nozzle, defined CS/IS/AL combinations (see Fig. 1E) yielded internal diameters from ∼150 to 300 µm, while the external diameter remained stable near 410 µm, as expected from mass conservation. With a 200 µm nozzle, corresponding combinations (see Fig. 1F) produced internal diameters of ∼50–150 µm, with external diameters around 210 µm. Notably, these size ranges closely correspond to those of lymphatic capillaries and collecting vessels observed in vivo [34–36].

Overall, the microfluidic extrusion consistently produces size-controlled hydrogel tubes that support the self-organization of lymphatic endothelial cells into a circumferential monolayer lining a central lumen within seven days.

### 2. Matrix composition directs cellular behavior, drives extracellular remodeling, and supports structural maturation of 3D lymphatic endothelium

Based on prior studies highlighting the roles of gelatin (G), Matrigel (MG), hyaluronic acid (HA), or fibrinogen (FG) in endothelial network formation and extracellular matrix (ECM) remodeling [32, 37–41], we tested 9 matrix formulations. We combined one to four of these components at fixed total concentrations (w/v) to investigate potential synergistic effects (Fig. 1A and Table 1), varying the specific identity and combination of constituents. To ensure precisely defined and reproducible experimental conditions, we employed natural proteins of well-characterized composition together with growth factor-reduced Matrigel.

**Table 1.** Morphological features of lymphatic cell organization within tubular alginate constructs across different ECM formulations after 7 days of culture Formulations combined natural ECM-derived proteins—gelatin, Matrigel, hyaluronic acid, and fibrinogen at fixed concentrations (in w/v), with variation limited to component identity. Lymphatic endothelial cells were encapsulated at 10□ cells/mL. To focus on the role of ECM identity, concentrations were fixed using mid-range values widely established in the literature, avoiding confounding effects from dose variability.

| Formulation | % Composition (in w/v) | Day 7 – Observed Outcome |
| --- | --- | --- |
| Gelatin (G) | 2% | Sparse, round, and isolated cells |
| Matrigel (MG) | 30% | Cord-like structures extending toward the alginate wall |
| G/Hyaluronic Acid (HA) | 2%/0.2% | Central aggregates of rounded cells; no axial alignment |
| MG/HA | 30%/0.2% | Small rounded aggregates near the alginate wall |
| G/FG (Fibrinogen) | 2%/0.2% | Elongated cords centrally aligned along the tube axis |
| MG/FG | 30%/0.2% | Dense cords along the alginate wall; incomplete surface coverage |
| G/HA/FG | 2%/0.2%/0.2% | Irregular linear cords with moderate continuity |
| MG/HA/FG | 30%/0.2%/0.2% | Hollow tubular structures; continuous and elongated |
| G/MG/HA/FG | 2%/30%/0.2%/0.2% | Continuous, elongated, and well-organized hollow tubular structures |

The observed cellular outcomes, reported in Table 1, reveal that only two formulations, namely MG/HA/FG and G/MG/HA/FG, supported the formation of vessel-like architectures. In contrast, single-component matrices (G or MG) yielded sparsely distributed or rounded cells, while two-component combinations generally failed to induce lumen formation. HA-containing mixtures preserved rounded cell morphology, whereas FG-containing combinations promoted partial nucleation of cellular structures. The G/HA/FG mixture exhibited intermediate behavior, with cells partially assembled into cord-like structures but lacking cohesive or fully organized architecture. Furthermore, to study self-assembly and matrix remodeling, we compared extruded hydrogel tubes with alginate-free droplet cultures using 12 formulations of four matrix components over seven days. Supplementary Tables S1 and S2 provide morphological data on lymphatic cell organization across these formulations. Only MG/HA/FG and G/MG/HA/FG consistently formed compact, cord-like structures in alginate-free droplets and continuous, hollow monolayers in extruded tubes.

To investigate the cellular architecture associated to structural maturation, we performed immunofluorescence staining experiments with confocal microscopy at day 14 on MG/HA/FG and G/MG/HA/FG constructs - the only formulations yielding vessel-like structures. MG/HA/FG samples exhibited fragmented actin networks and dispersed nuclei (Fig. 2B), whereas G/MG/HA/FG constructs displayed continuous monolayers with uniform nuclear distribution and cortical actin (Fig. 2C). To analyze actin organization, images were first enhanced using top-hat filtering to improve filament contrast, then binarized to isolate actin structures. From these binarized images, radial autocorrelation profiles were extracted (Supplementary Fig. S5B), revealing a steeper decay in MG/HA/FG samples compared to G/MG/HA/FG, indicative of greater spatial disorder. Further spatial analysis based on distance transformation [42] (Fig. 2E) and nearest-neighbor distance histograms (Fig. 2F) showed increased spacing between actin structures in MG/HA/FG constructs relative to G/MG/HA/FG (16 µm vs 5 µm in mean inter-distance), indicative of a more disorganized actin distribution. In addition, local entropy measurements (Supplementary Fig. S5D) confirmed a higher degree of spatial disorder in MG/HA/FG, collectively reflecting a more organized cytoskeletal network in the four-component matrix.

**Figure 2.**
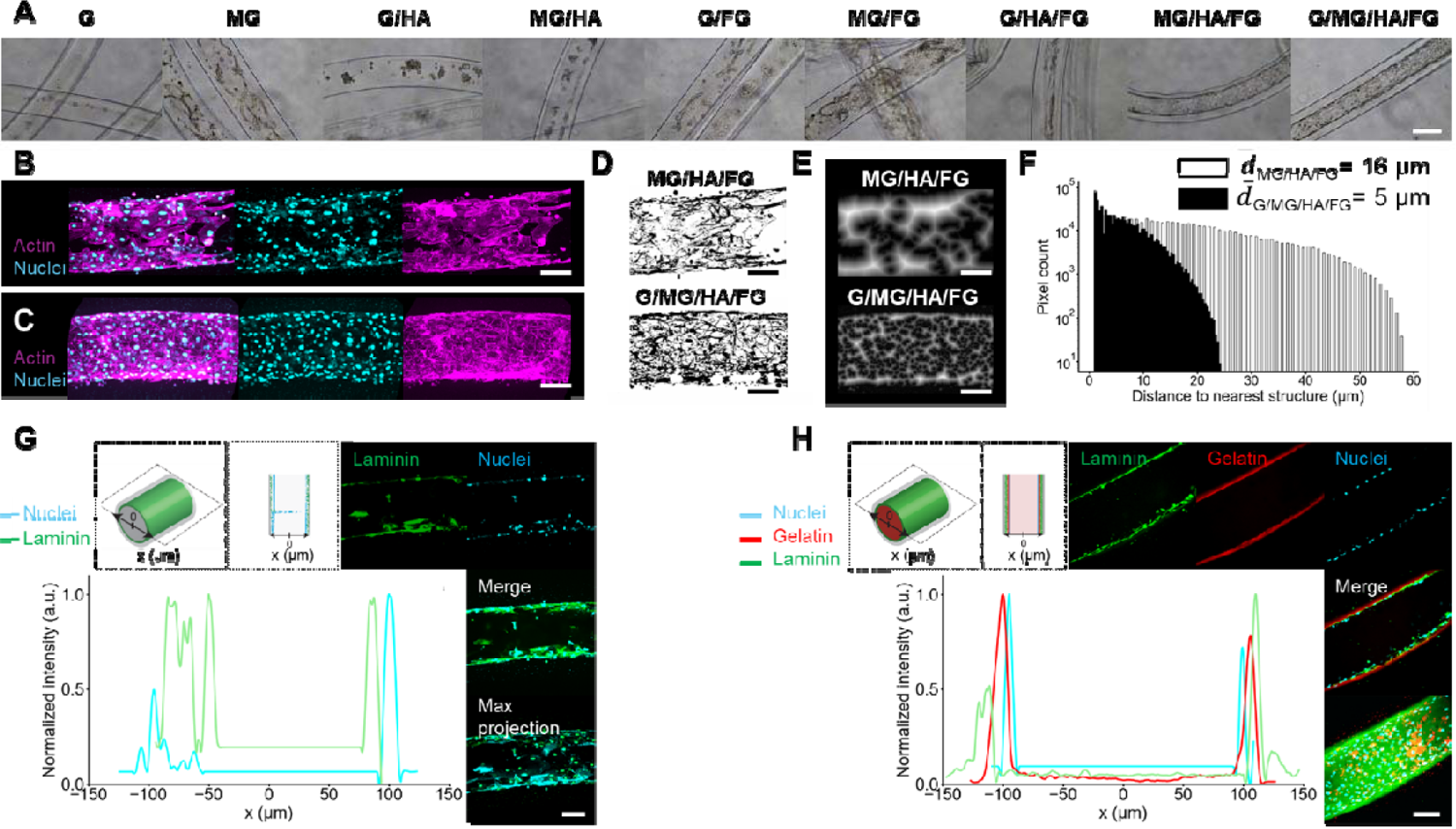
Matrix formulation as a critical factor in HDLEC self-organization, lumen formation, and progressive matrix remodeling in *in vitro* 3D lymphatic tubes. **(A)** Brightfield images of constructs at day 7 fabricated with 1-, 2-, 3-, and 4-component matrices. Only MG/HA/FG and G/MG/HA/FG formulations supported vessel-like tube formation at low HDLEC seeding density. **(B–C)** Confocal maximum intensity projections of day-14 tubes stained for actin (magenta) and nuclei (cyan). MG/HA/FG constructs **(B)** display fragmented cytoskeletal networks and uneven nuclear distribution. In contrast, G/MG/HA/FG constructs **(C)** exhibit continuous cortical actin and uniformly distributed nuclei consistent with monolayer formation. **(D)** Top-hat filtered binarized images derived from confocal actin filament images of entire representative tubes. **(E)** Euclidean distance transform (EDT) maps generated from the processed confocal actin images shown in panel **D**. **(F)** Histogram of nearest-neighbor distances quantifying the spatial organization of actin structures. Values were derived from Euclidean distance transform (EDT) maps (see panel **E**) for the two conditions: G/MG/HA/FG (solid black bars) and MG/HA/FG (white bars with black outlines). The distributions are plotted on a logarithmic scale and the mean spacing values 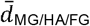 and 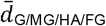 are indicated accordingly. **(G)** Confocal cross-section of a MG/HA/FG construct at day 14, showing a 3D view of the equatorial plane of an alginate tube alongside a schematic 2D equatorial representation (laminin: cyan; nuclei: green). Individual fluorescence images of laminin, nuclei, and merged signals are presented, with corresponding fluorescence intensity profiles plotted along the x-axis. **(H)** Confocal cross-section of a G/MG/HA/FG construct at day 14, showing a schematic 2D equatorial representation (laminin: cyan; gelatin: red; nuclei: green) followed by fluorescence images of laminin, nuclei, gelatin, and merged composite. Laminin localizes at the outer alginate interface, gelatin occupies the intermediate zone, and HDLECs form a continuous inner lining. Intensity profiles along the x-axis confirm spatial segregation of laminin, nuclei, and gelatin, indicating the formation of a stratified tubular architecture. **Scale bars:** 100 µm **(A–E)**, 50 µm **(F–G)**.

To elucidate the structural differences between MG/HA/FG and G/MG/HA/FG matrices, we examined the spatial distribution of key matrix components—laminin (the principal constituent of Matrigel) and gelatin—as well as the distribution profiles of cell nuclei at day 14. In MG/HA/FG constructs, HDLECs and laminin exhibited partial alignment along the core–alginate interface; however, their distribution remained scattered and disorganized, with no discernible spatial layering, suggestive of progressive tissue architecture collapse at this time point (Fig. 2G). By contrast, G/MG/HA/FG constructs displayed a well-defined organization: HDLECs formed a continuous inner lining, with laminin concentrated at the alginate wall, and gelatin sandwiched in between (Fig. 2H). To assess the spatial organization, we compared cell-laden constructs to those without HDLECs at days 1 and 14 (Fig. S6–S7). In the absence of cells, laminin passively adsorbed to the alginate boundary, and gelatin maintained an overall uniform distribution within the core, lacking clear localization patterns. In contrast, HDLEC-seeded constructs showed progressive confinement of gelatin to the intermediate zone, consistent with cell-mediated matrix restructuring over time.

These findings indicate that the G/MG/HA/FG matrix not only facilitates early cellular organization but also supports ongoing matrix remodeling, which contributes to the formation of structurally organized lymphatic endothelium.

### 3. Long-term proliferative dynamics, viability, and homeostasis in 3D lymphatic endothelium

Building on the early cellular organization and matrix remodeling seen in the G/MG/HA/FG construct, we next assessed the long-term behavior of the 3D lymphatic endothelium. We focused on proliferation, viability, and structural. Mitotic activity, assessed by phospho-histone H3 (PHH3) staining, was high at day 5 (52% PHH3-positive cells), sharply declined by day 14 (9%), and was nearly absent by day 30 (3%) (Fig. 3A–B) [43]. In parallel, Ki-67 staining mirrored the PHH3 profile, with elevated early expression followed by a gradual reduction by day 30, consistent with decreased proliferative activity over time (Fig. S10). Furthermore, viability analysis showed that the percentage of dead cells remained stable between day 14 (7%) and day 30 (7%) (Fig. 3C).

**Figure 3.**
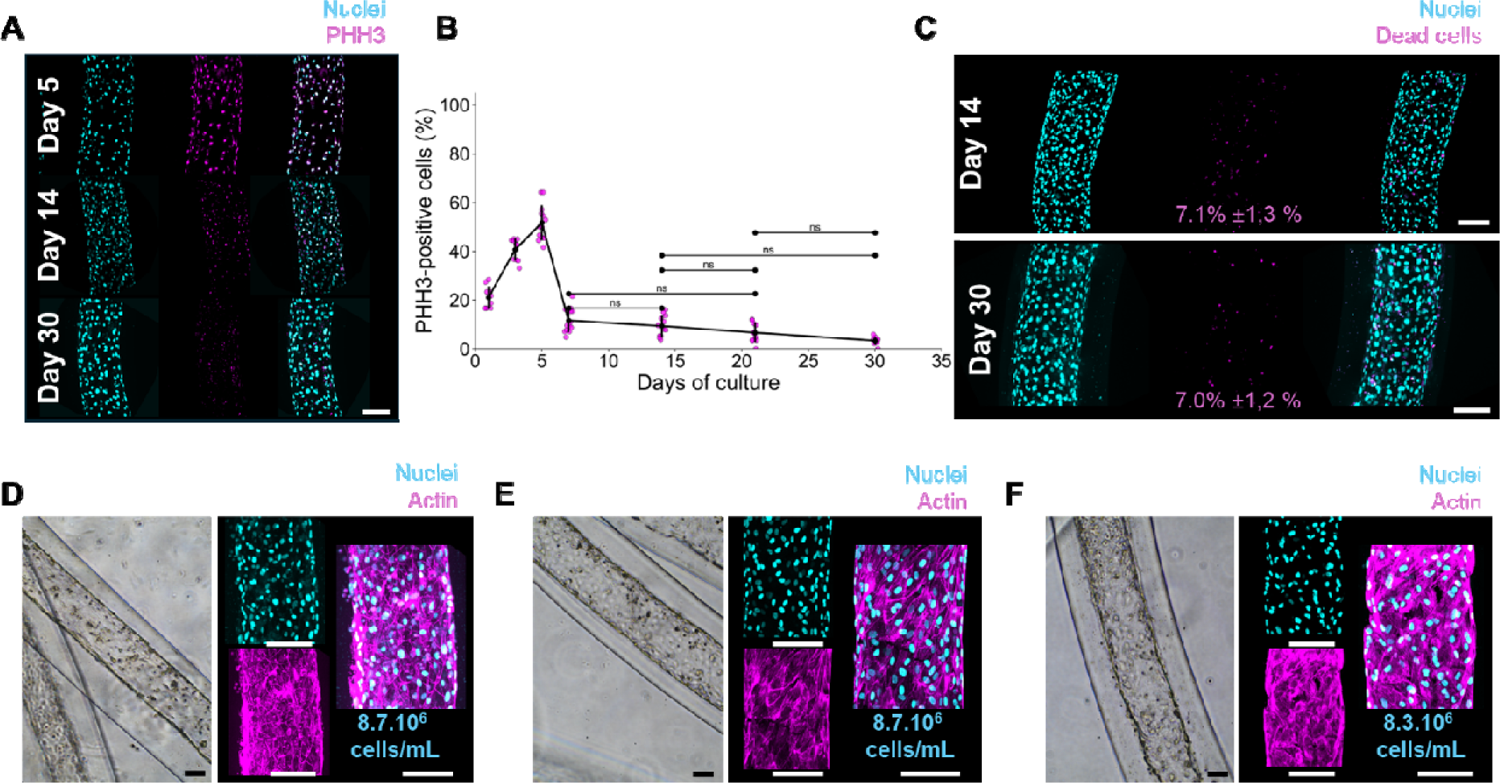
Long-term viability and structural stability of *in vitro* 3D lymphatic endothelium. **(A)** Maximal projections showing nuclei (cyan) and phospho-histone H3 (PHH3, magenta), along with merged channels, at days 5, 14, and 30, illustrating the decline in mitotic activity. **(B)** Dot plot quantifying the percentage of PHH3-positive cells relative to total nuclei, demonstrating a mitotic index greater than 50% at day 5, less than 10% at day 14, and below 3.5% at day 30. (N≥ 3 constructs per condition. **(C)** Maximal projections showing nuclei (cyan) and dead cells (magenta), with merged channels. Quantification of viability based on nuclear/viability staining (central merged panel) indicates low and stable mortality over time (day 14: 7.1 ± 1.3%; day 30: 7.0 ± 1.2%, N≥3 constructs per condition). **(D–F)** Brightfield and confocal maximum intensity projections of HDLEC constructs at days 14 (**D)**, 21 **(E)**, and 30 **(F)**, showing a dense and preserved HDLEC monolayer. Nuclei (cyan) and actin (magenta) remain evenly distributed with cortical enrichment. **Scale bars:** 100 µm.

To assess morphological stability, we performed a temporal analysis from day 14 to day 30. Confocal imaging of longitudinal sections revealed preservation of the cylindrical architecture and an intact luminal lining at days 14, 21, and 30 (Fig. 3D). The cortical actin signal remained continuous, and nuclei were evenly spaced along the tube periphery, suggesting sustained monolayer integrity and structural preservation (Fig. 3E–F). These observations were further supported by nuclear quantification on maximum intensity projections from transverse optical sections (Fig. 3D–F), which showed stable cell densities ranging from 8.7 × 10□ cells/mL at days 14 and 21 to 8.3 × 10□ cells/mL at day 30.

Matrix remodeling was assessed by monitoring the degradation of rhodamine-labeled gelatin from day 0 to day 28. A progressive loss of signal was observed, with near-complete depletion by day 13, except for a faint residual lining at the alginate interface (Fig. S8, Movies S1-S2). Endpoint confocal imaging at day 30 confirmed that HDLEC-laden constructs maintained the stratified matrix organization observed at day 14, characterized by nuclei lining the lumen, gelatin localized centrally, and laminin concentrated near the alginate wall (Fig. 2G, Movie S2, and Fig. S9).

To further examine endothelial behavior beyond day 30, we analyzed constructs at days 40 and 50. Confocal imaging confirmed sustained cortical actin organization and uniform nuclear distribution (Fig. S11A, B, D, E). Viability assays showed consistently low cell death (8%) at both time points (Fig. S11C, F). Ki-67 staining revealed low proliferative activity at days 40 and 50 (6 % and 7.5%, respectively; Fig. S11G). This phenomenon mirrors the dynamic equilibrium found in native lymphatic tissues, where both structural and functional stability are preserved despite ongoing molecular and cellular turnover [4, 44].

Together, these findings indicate that the 3D lymphatic endothelium formed within the G/MG/HA/FG matrix reaches homeostasis (low proliferative activity, limited yet sustained cell death, stable cell density, and preserved structural integrity). The latter are key hallmarks of mature lymphatic tissues in vivo [4, 45, 46].

### 4. Blood endothelial cells self-organize into long-lasting 3D monolayers under the same matrix conditions

To further explore how endothelial identity influences self-organization and function in 3D matrices, we compared HDLECs with two blood vascular endothelial lineages, human dermal microvascular endothelial cells (HDMECs) and human umbilical vein endothelial cells (HUVECs). All three cell types were cultured within the optimized G/MG/HA/FG hydrogel, and their morphological, molecular, and functional features were assessed over time.

Brightfield imaging of HDMEC- and HUVEC-based constructs showed that the cylindrical geometry was preserved from day 7 through day 30 (Fig. 4A, 4C). The inner tube walls progressively darkened, particularly by day 30, suggesting increased cell layer thickness.

**Figure 4.**
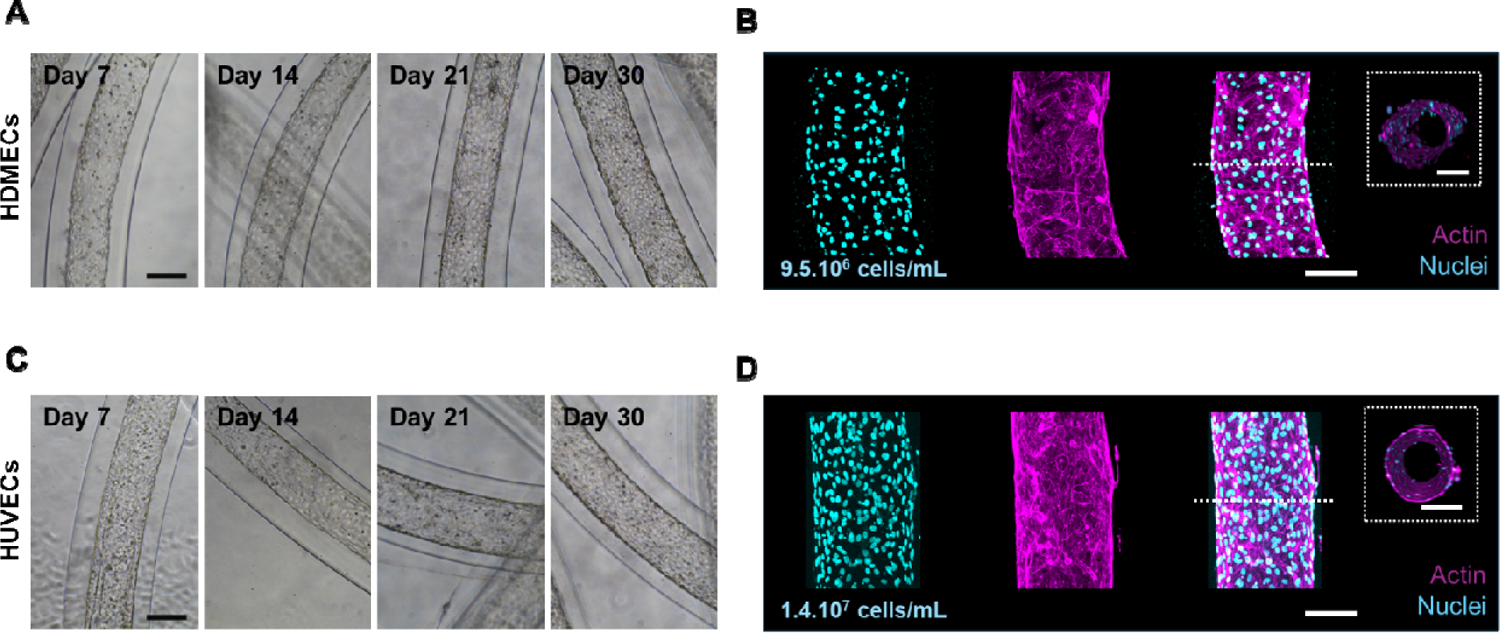
Sustained maintenance of lineage-specific 3D endothelial monolayers in the optimized hydrogel matrix. **(A, C)** Brightfield images of HDMEC-based **(A)** and HUVEC-based **(C)** constructs at days 7, 14, 21, and 30 of culture. Both cell types maintain a dense and stable monolayer over time. The progressive darkening of the tube walls suggests increased cellular compaction compared to HDLEC-based structures (see **Fig. 3A**). **(B, D)** Confocal maximum intensity projections of HDMEC-based **(B)** and HUVEC-based **(D)** constructs at day 30, showing nuclei (cyan), actin (magenta), and merged images. Insets show 3D reconstructions with selected viewing angles, confirming the formation of continuous monolayers fully covering the luminal surface. Nuclear density is higher than in HDLEC-based constructs, with approximately 9.6.10^6^ cells/mL for HDMECs and 1.5.10^7^ cells/mL for HUVECs. N≥3 independent extrusion batches were performed per condition **Scale bars:** 100 µm.

Confocal imaging at day 30 revealed that both HDMECs (Fig. 4B) and HUVECs (Fig. 4D) formed continuous luminal monolayers enriched with cortical actin and regularly spaced nuclei. Transverse sections confirmed complete coverage and lumen formation. Cell density was found to be 9.5×10^6^ cells/mL for HDMECs and 14×10^6^ cells/mL for HUVECs —both exceeding those observed in HDLEC constructs (Fig. 3B). However, no multilayering or architectural disruption was detected. At this point, the differences in cell densities between HDMEC, HUVEC and HDLEC constructs may reflect inherent variations in junctional properties specific to each endothelial lineage. This interpretation is consistent with established literature describing distinct junctional characteristics between blood and lymphatic endothelia [47, 48], though further studies are required to confirm this finding in our model.

### 5. Engineered lymphatic and blood endothelial monolayers exhibit lineage-specific features in structure, markers, and barrier function

To further characterize each endothelial lineage, we analyzed marker expression and barrier properties at day 30. Confocal imaging of HDLEC-based constructs (Fig. 5A) revealed a dense, cohesive monolayer enriched with cortical actin and strong expression of CD31, LYVE-1, and podoplanin (PDPN), confirming sustained expression of lymphatic-specific markers. After 30 days of culture, fibronectin (FN) predominantly localized to the basal compartment, indicating active extracellular matrix production by HDLECs (see also Fig. S12A for day 14). As expected, HDMEC- and HUVEC-derived monolayers exhibited similarly organized cell layers with strong CD31 expression (Fig. 5B–C; Fig. S12B–C). Furthermore, HDMECs lacked both LYVE-1 and PDPN expressions, whereas HUVECs displayed a LYVE-1 signal but no detectable PDPN, the latter consistent with their respective blood endothelial identity.

**Figure 5.**
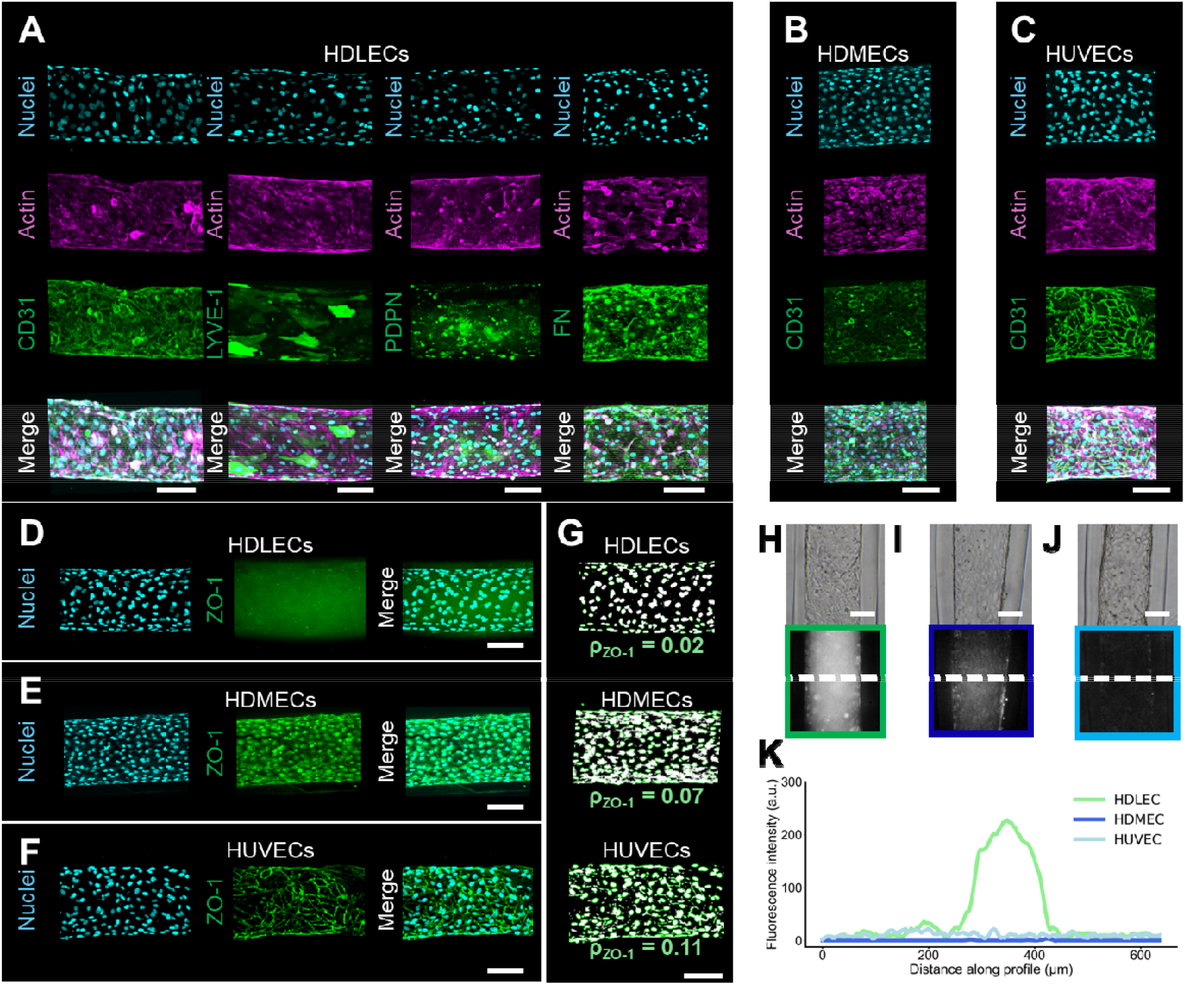
Lineage-specific molecular features and barrier function in in vitro 3D endothelial monolayers. **(A–C)** Confocal maximum intensity projections of HDLEC-, HDMEC-, and HUVEC-based constructs. Each row presents individual channels for nuclei (cyan), actin (magenta), and various lineage-specific markers (green), followed by merged images. **(A)** HDLECs express CD31 (pan-endothelial marker), LYVE-1 and podoplanin (PDPN) (lymphatic-specific markers), and fibronectin (FN), which is secreted by endothelial cells. **(B–C)** HDMECs **(B)** and HUVECs **(C)** express CD31 consistent with their blood endothelial identity. **(D–F)** Confocal maximum intensity projections of ZO-1 immunostaining at day 30. **(D)** ZO-1 signal in HDLEC constructs is weak, fragmented, and mostly indistinct from background, consistent with poorly organized junctions. **(E–F)** HDMEC **(E)** and HUVEC **(F)** constructs display stronger, continuous junctional staining. Merged views include nuclei (cyan). **(G)** Binary segmentation masks of ZO-1 staining. ZO-1 density (ρ_ZO-1_) is defined as the percentage of image area occupied by ZO-1-positive pixels. HDLEC constructs exhibit sparse, fragmented junctions (ρ_ZO-1_ = 0.02), whereas HDMECs (ρ_ZO-1_ = 0.07) and HUVECs (ρ_ZO-1_ = 0.11) form denser, continuous junctional networks. **(H–J)** Fluorescence images following incubation with 70 kDa rhodamine-labeled dextran. **(H)** Only HDLEC constructs allow tracer accumulation in the lumen. **(I–J)** HDMEC and HUVEC constructs largely exclude the dextran. Insets show higher magnification views of the lumen. **(K)** Quantification of luminal dextran fluorescence intensity across the lumen midsection (x-axis: transverse position in µm; y-axis: fluorescence intensity in arbitrary units). HDLEC constructs (green) exhibit a clear central peak, indicating luminal tracer entry, whereas HDMEC (royal blue) and HUVEC (light blue) profiles remain flat, confirming exclusion. **Scale bars:** 100 µm.

We next investigated the organization of intercellular junctions, a key structural feature of endothelial barriers. Given their functional role in regulating permeability, we focused on tight junctions and analyzed ZO-1 distribution as a representative marker. In HDLEC-derived tubes (Fig. 5D), ZO-1 staining appeared faint and discontinuous— consistent with the loosely organized junctions typical of lymphatic endothelium. In contrast, HDMECs (Fig. 5E) and HUVECs (Fig. 5F) displayed strong, continuous ZO-1 labeling along cell–cell interfaces, reflecting the tight intercellular contacts characteristic of blood endothelial layers. Notably, all three cell types expressed ZO-1 in 2D monolayers (Fig. S13), indicating that the differences observed in 3D arise from spatial reorganization within the tubular environment rather than from loss of expression.

To quantitatively compare junctional architecture across endothelial lineages, we carried out a semi-quantitative analysis of ZO-1 distribution. Using an image processing pipeline involving Gaussian smoothing, Sobel-based edge enhancement, and adaptive thresholding, we generated binary masks of ZO-1-positive regions (Fig. 5G). The 3D endothelium formed by HDLECs showed sparse and fragmented ZO-1 labeling (density ρ_ZO-1_ = 0.02), sharply contrasting with HDMECs (ρ_ZO-1_ = 0.07) and HUVECs (ρ_ZO-1_ = 0.11), which exhibited more extensive and cohesive intercellular junctions.

To assess macromolecular retention within the lumen, we exposed tubes to a 70 kDa rhodamine-labeled dextran tracer while excluding the tube ends from the fluorescent solution (Fig. S14A). This setup confined the tracer to the surrounding medium, requiring diffusion through the outer alginate wall and the basal endothelial surface to reach the lumen, allowing direct measurement of transendothelial permeability. Only HDLEC-derived endothelium (Fig. 5I) showed detectable luminal tracer accumulation, indicating successful passage of the 70 kDa dextran. By contrast, HDMEC- and HUVEC-derived endothelia (Fig. 5J–K) displayed no intraluminal signal, consistent with a more restrictive barrier. These findings were confirmed by low-magnification fluorescence imaging (Fig. S14B–D), where tracer accumulation remained clearly visible in HDLEC lumen, while no signal was detected in HDMEC or HUVEC constructs. Fluorescence intensity profiles across the lumen (Fig. S14E) confirmed these findings: only HDLEC endothelium exhibited a distinct intraluminal peak (green trace), whereas HDMEC (dark blue) and HUVEC (light blue) profiles remained flat.

To evaluate size selectivity, we repeated the permeability assay using dextran molecules of different molecular weights. The 3 kDa tracer freely diffused into the lumen of all constructs, indicating unrestricted passage of small solutes across each endothelial barrier. The 500 kDa tracer was excluded from all lumens, irrespective of cell type (Fig. S15). Notably, 3 kDa and 70 kDa dextrans are known to permeate the alginate-based matrices used here, while 500 kDa dextrans are effectively retained by the gel [49, 50]. Therefore, only the 3 kDa and 70 kDa tracers were appropriate for probing differences in endothelial permeability. Within this range, the selective accumulation of 70 kDa dextran in HDLEC-derived tubes may reflect intrinsic properties of the endothelial layer—specifically, its permissiveness to macromolecular transport, a hallmark of lymphatic capillaries [51, 52].

Altogether, these findings demonstrate that HDLEC-, HDMEC-, and HUVEC-derived endothelia maintain distinct and stable characteristics over 30 days of 3D culture under identical matrix conditions. Each lineage exhibited a consistent molecular profile, well-defined junctional architecture, and characteristic permeability behavior. Notably, HDLEC-derived monolayers preserved lymphatic-specific marker expression, showed sparse and discontinuous ZO-1-positive junctions, and selectively permitted macromolecular passage into the lumen—key hallmarks of lymphatic endothelial identity. Beyond supporting lymphatic structures, this platform also enables the formation of stable, long-lasting 3D blood endothelial layers, providing a valuable benchmark for comparative studies. The convergence of structural, molecular, and functional features underscores the robustness of this system in maintaining lymphatic endothelial identity within a defined 3D environment.

## Discussion

This study reports a robust, tunable, and scalable platform for engineering a 3D lymphatic endothelium. We used a microfluidic extrusion-based technique and optimized a protein matrix mixture. Using primary HDLECs, we show that the model overcomes limitations of immortalized or animal-derived models, while preserving cellular identity. The constructs recapitulate key hallmarks of lymphatic vessels, i.e. monolayer architecture, selective marker expression, and permeability. They also allow precise size control and provide sustained long-term maintenance.

We observe that HDLECs seeded within tubular constructs monolayers self-organized around a cylindrical lumen over 7 days. This process mimics early lymphatic assembly *in vivo*, where dispersed endothelial progenitors align and form lumenized structures [53, 54]. Importantly, although culture is performed under static conditions, the *in vitro* monolayers remain structurally and cellularly stable for at least 30 days without needing perfusion or matrix renewal surpassing the typical lifespan of most 3D endothelial models, which often degrade or delaminate within less two weeks [51, 55, 56]. While perfusion is not required in lymphatic vessels generation, it is often considered to be highly beneficial to prevent lumen collapse and enhance long term functional relevance [57]. Our findings may suggest that flow/perfusion effects are counterbalanced by biochemical cues from the custom ECM mixture. This long-term stability is crucial in pharmacokinetics, or to study interactions with immune cells.

Matrix composition was systematically investigated to identify conditions supporting both early cellular organization and long-term structural maturation. Among all tested formulations, only the four-component G/MG/HA/FG matrix consistently enabled initial monolayer formation and its sustained structural refinement. Each component contributed distinct biophysical or biochemical cues: Fibrinogen enhanced adhesion and initial network formation [41]; Hyaluronic acid appeared to support the maintenance of nucleated cellular architectures, potentially synergizing with fibrinogen [32, 40]; Matrigel provided basement membrane proteins such as laminin and collagen IV [38, 39]; and gelatin promoted long-term cohesion and facilitated ECM remodeling [37]. This four component matrix represents a significant improvement of previous methods where MG was only used for vascular constructs [39]. The build-up of fibronectin and the formation of a layered structure indicates that HDLECs actively remodel the matrix, mirroring processes in lymphatic development and matrix regulation [58, 59].

Beyond maintaining structural integrity, the model preserves molecular and transport characteristics of lymphatic endothelial cells. HDLECs sustain expression of LYVE-1 and podoplanin and formed discontinuous junctions with sparse ZO-1 labeling, reflecting their role in macromolecular uptake. Accordingly, these vessel constructs permit luminal entry of 70 kDa dextran. In contrast, blood endothelial cell-lined constructs (HDMECs and HUVECs) exhibit continuous junctions, exclude the tracer, and lack lymphatic markers, underscoring lineage-specific differences under identical culture conditions [44, 45, 49]. Moreover, building on these observations and including additional morphological features such as physiological tube diameters and reduced cell density, HDLEC constructs collectively exhibit hallmarks of initial lymphatic capillaries (e.g. selective marker expression, loose intercellular junctions, and macromolecule permeability). The lymphatic endothelium has two distinct features related to intercellular junctions. First, lymphatics exhibit buttons which are characterized by fragmented intercellular endothelial junctions but also classical tight junctions which are found in endothelial cells from collecting lymphatic vessels [47]. The former are more permeable than the latter. Lymphatic endothelial cells used in our work are derived from skin tissue and more likely possess buttons and are thus more permeable. The absence of mural cells or valve-like structures further distinguishes these constructs from models of collecting vessels. Thus, our vessel constructs are more likely to mirror the situation of endothelial cells of initial lymphatic vessels.

An additional further step is the development of models for collecting lymphatics which incorporate additional cell types and have valve-forming capacity. Such advancements would enable more accurate modeling of lymphatic function and disease, broadening the platform’s utility across a range of biomedical applications.

The extrusion-based approach demonstrates its effectiveness with endothelial cells derived from various sources. Lymphatic and vascular endothelial cells (HDLECs, HDMECs, and HUVECs) both formed stable 3D monolayers exhibiting appropriate spatial organization. They differ in cell density, junctional continuity, and permeability that are due to lineage-specific traits rather than to matrix-related cues.

In summary, the integration of extrusion technology with a finely tuned hydrogel matrix offers several key benefits: it allows for precise shaping of vessel constructs, ensures consistent and long-term culture conditions, and supports a variety of endothelial cell types. This approach provides a reliable method for developing three-dimensional endothelial vessels that faithfully retain the unique identity and functions of different endothelial lineages. Beyond its use in modeling lymphatic vessels, the platform stands as a powerful tool for comparative vascular studies, drug transport research, and investigations into endothelial adaptability and disease-associated alterations. Its robust, modular nature makes it highly adaptable, paving the way for the study of complex vascular biology and the development of novel therapeutic strategies.

## Materials and methods

### 1. Cell culture

Primary human dermal lymphatic endothelial cells (HDLECs; PromoCell, C-12216) and human dermal microvascular endothelial cells (HDMECs; PromoCell, C-12212) were cultured in Endothelial Cell Growth Medium MV2 (EGM2-MV; PromoCell, C-22022). Human umbilical vein endothelial cells (HUVECs; Lonza, CC-2519) were cultured in Endothelial Cell Growth Medium 2 (EGM2; PromoCell, C-22011).

Penicillin–streptomycin (1%, Gibco, 15140122) was included in all cultures. Cells were maintained in tissue culture-treated flasks at 37 °C in a humidified 5% CO_2_ incubator and used between passages 2 and 6. Before encapsulation, cells were detached using 0.05% trypsin–EDTA (Gibco, 25300054), centrifuged at 400 × *g* for 5 minutes, and resuspended in hydrogel matrices at a final density of 10□ cells/mL unless otherwise specified.

### 2. Hydrogel formulation

Hydrogel matrices were prepared using combinations of gelatin (2% w/v; porcine, Type A, Advanced BioMatrix, 5005), growth factor-reduced Matrigel (30% w/v; Corning, 356231), low molecular weight hyaluronic acid (100–150 kDa, 0.2% w/v; Advanced BioMatrix, 5003), and human fibrinogen (0.2% w/v; Sigma-Aldrich, F8630). All components except gelatin were kept at 4 °C; gelatin was added just before encapsulation to avoid premature gelling.

Nine matrix formulations were prepared by combining one to four macromolecular components—gelatin (G; 2% w/v), Matrigel (MG; 30% w/v), hyaluronic acid (HA; 0.2% w/v), and fibrinogen (FG; 0.2% w/v). Each component was used at a fixed final concentration when included, regardless of whether it was part of a single-, double-, triple-, or four-component mixture. Acellular matrices were processed identically for control experiments.

### 3. Coaxial extrusion of tubular constructs

The coaxial extrusion setup was adapted from previous designs for encapsulated hydrogel structures [35, 53, 54]. A custom triple coaxial head was 3D-printed using an EnvisionTEC D4K Pro printer and HTM 140 V2 resin. Three syringe pumps (neMESYS low pressure module, CETONI) controlled flow rates independently.

The core stream contained the cell–hydrogel mix at 4–7 °C, with cells added at 10□ cells/mL. The intermediate stream was 0.6 M sorbitol (Sigma-Aldrich, S1876), and the shell stream was 2% sodium alginate (Alliance Gums & Industries, I1G80-E401), sterilized by autoclaving.

Extrusion occurred into a 100 mM CaCl_2_ bath (37 °C) to rapidly gel the alginate. Flow rates were typically 2.0□mL/h (shell), 1.0□mL/h (intermediate), and 1.0□mL/h (core) unless otherwise stated. Two nozzle sizes (450 µm and 200 µm) were used to vary tube diameter over an extended range. Within 10 minutes post-extrusion, tubes were transferred to 60□mm Petri dishes (Corning, CLS430166) containing 5□mL of warm culture medium and incubated at 37 °C, 5% CO_2_.

### 4. Culture conditions and experimental time course

After extrusion, constructs were cultured in 60□mm Petri dishes (Corning) with 5□mL of endothelial medium per dish. HDLEC and HDMEC constructs were maintained in EGM2-MV medium (PromoCell, C-22022), while HUVEC constructs received EGM2 (PromoCell, C-22011). Media were refreshed every 48□hours by gentle aspiration and replacement. No perfusion or mechanical agitation was applied throughout the culture period.

Constructs were monitored and collected at specific time points: day 0, 1, 2, 3, 5, 7, 14, 21, and 30. Optional late-stage endpoints at day 40 and 50 were included in selected experiments. At each time point, constructs were either processed for viability analysis, barrier permeability assays, or quantitative image acquisition or fixed for immunostaining.

### 5. Cell viability assays

Cell viability was at selected time points (days 7, 14, 21, 30, 40, and 50) using the LIVE/DEAD Viability/Cytotoxicity Kit (Thermo Fisher Scientific, L3224), following the manufacturer’s protocol. Briefly, 1-cm segments of hydrogel constructs were carefully transferred into 60□mm Petri dishes (Corning, CLS430166) pre-coated with poly-L-lysine (PLL; Gibco, A3890401) to enhance construct adhesion and minimize movement during staining and imaging. Samples were incubated for 30 minutes at 37 °C in a staining solution containing 2 µM calcein-AM and 4 µM ethidium homodimer-1 diluted in phenol red–free DMEM. After a gentle rinse in fresh medium, constructs were immediately imaged using the spinning-disk confocal microscope described in Section 6.

### 6. Immunofluorescence staining

At designated time points, 1-cm segments of hydrogel constructs were carefully transferred to 1.5□mL microtubes and fixed in 4% paraformaldehyde (PFA; Fisher Scientific, F/1501/PB17) diluted in Dulbecco’s modified Eagle’s medium (DMEM; PAN-Biotech, P04-05545) for 1□hour at room temperature under gentle agitation. Samples were then washed three times with DMEM and stored at 4 °C until further processing.

For staining, samples were permeabilized for 30□minutes in DMEM with 1% Triton X-100 (Sigma-Aldrich, T8787), then blocked for 1□hour in DMEM containing 1% bovine serum albumin (BSA; Sigma-Aldrich, A9647) and 10% fetal bovine serum (FBS; Clinisciences, FBS Xtra, FBS-16A) to reduce nonspecific binding. All steps were performed under gentle orbital shaking (40□rpm) to facilitate diffusion into the tubular lumen.

Constructs were incubated overnight at 4 °C with primary antibodies diluted in blocking buffer supplemented with 0.1% Triton X-100 and 2% FBS. The following primary antibodies were used:

- CD31 (Polyclonal Sheep IgG, 1:200; BioTechne, AF806),
- LYVE-1 (Rabbit monoclonal, 1:500; Abcam, ab219556),
- Podoplanin /PDPN (Mouse monoclonal, 1:500; MBL Life Science, 8F11),
- Fibronectin (Rabbit polyclonal, 1:500; Abcam, ab2413),
- ZO-1 (Mouse monoclonal, 1:500; Thermo Fisher Scientific, ZO1-1A12),
- Phospho-Histone H3 (Rabbit polyclonal, 1:500; Sigma-Aldrich, SAB4502231),
- Ki-67 (Rabbit polyclonal, 1:500; Abcam, ab15580),
- Laminin (Rabbit polyclonal, 1:200; Abcam, ab11575).

After DMEM washes, samples were incubated overnight at 4 °C with appropriate secondary antibodies conjugated to Alexa Fluor 488, 568, or 647 (1:500; Thermo Fisher Scientific). Actin filaments were visualized using Phalloidin–Alexa Fluor 568 (1:200; Thermo Fisher Scientific, A12380), and nuclei were counterstained with Hoechst 33342 (1 µg/mL; Thermo Fisher Scientific, H3570).

Following staining, samples were optically cleared by sequential immersion in 30%, 60%, and 100% FUnGI solution to enhance imaging quality. FUnGI is a clearing agent composed of 50% glycerol (w/v), 2.5□M fructose, 2.5□M urea, 10.6□mM Tris base, and 1□mM EDTA, prepared as described in [60]. This graded clearing protocol was optimized to preserve fluorescence while progressively matching the refractive index of the tissue.

Cleared constructs were mounted between glass slides (RS France, BP024) using iSpacer chambers (0.25□mm and 0.15□mm; SunJin Lab, IS216 and IS015) and glass coverslips (Merck-Sigma, CLS2980223), then stored at 4 °C until imaging.

### 7. Brightfield and confocal imaging

The morphological evolution of constructs was first monitored in brightfield using a Nikon Eclipse Ti inverted microscope equipped with phase-contrast objectives and a digital camera (DS-Fi3, Nikon). Images were acquired directly from live constructs in culture dishes at predefined time points (days 0, 2, 5, 7, 14, 21, and 30). Acquisition parameters were kept constant across samples using NIS-Elements software (Nikon). Moreover, supplemental movies S1 and S2 were acquired using an *Incubascope*, a compact microscope inside a standard incubator, enabling transmitted light and fluorescence imaging over large fields of view directly inside a standard incubator, as described in [61].

In parallel, fixed and immunostained samples were imaged by spinning-disk confocal microscopy using a Leica DMI8 inverted microscope (Leica Microsystems, Wetzlar, Germany) equipped with a CSU-W1 spinning-disk scanner (Yokogawa Electric Corporation) and a LiveSR super-resolution module (Gataca Systems). The microscope was operated under environmental control (37 °C, 5% CO_2_) and controlled via MetaMorph software (Molecular Devices), using a 20× multi-immersion objective (Leica HC PL APO CS2, NA 0.75, working distance 0.66□mm).

Fluorophore excitation was performed using laser lines at 405, 488, 561, and 642□nm, and fluorescence emission was recorded with a Prime 95B sCMOS camera (Photometrics). Z-stacks were acquired with a step size of 0.55 µm and processed using FIJI (ImageJ) to generate maximum intensity projections. Brightness and contrast were adjusted uniformly across datasets. No deconvolution, denoising, or nonlinear filtering was applied.

### 8. Image analysis

Nuclei were segmented from maximum intensity projections in FIJI using Gaussian blur, Otsu thresholding, and watershed algorithms. Rectangular ROIs (200 µm × 500 µm) were defined to compute cell density (cells/mL), and nearest-neighbor distances were calculated to evaluate nuclear spacing and spatial organization. Phalloidin-stained actin filaments were processed by top-hat filtering followed by Phansalkar binarization. Euclidean Distance Transform (EDT) maps were generated to extract filament spacing, and peak-to-peak EDT distances were used to quantify cytoskeletal compactness.

Proliferating cells positive for phospho-Histone H3 (PHH3) or Ki-67 were quantified using the same segmentation workflow. DAPI-stained nuclei were first identified, and Otsu thresholding was then applied to the corresponding fluorescence channels to detect Ki-67^+^ or PHH3^+^ nuclei. Live (calcein^+^) and dead (ethidium^+^) cells were similarly segmented from z-stack projections. In both cases, marker-positive cells were isolated by watershed separation following thresholding. Parallel Python scripts replicating the same segmentation logic were used to validate the results. Quantification was expressed as the percentage of marker-positive cells relative to total nuclei per field.

ZO-1 immunostaining was analyzed on maximum projections after image smoothing, Sobel edge detection, and Otsu thresholding. Junctional density was defined as the percentage of the image area covered by ZO-1 signal. At least five fields per condition were analyzed using standardized acquisition and processing parameters.

### 9. Permeability assays

Permeability was tested using rhodamine B–conjugated dextrans: 3□kDa (Invitrogen, D3307), 70□kDa (Invitrogen, D1841), and 500□kDa (HAworks, DE-Rhodamine-500k). Tubes were incubated for 30 minutes at 37 °C in culture medium containing 100 µg/mL tracer. Both ends of the tube were kept fully outside the fluorescent solution to ensure passive transwall diffusion only. Tubes were handled carefully to preserve structural integrity.

After incubation, constructs were rinsed three times with fresh pre-warmed medium (EGM2 or EGM2-MV, depending on cell type). Fluorescence and brightfield images were acquired using the Nikon Eclipse Ti microscope under identical settings. Intensity profiles across the lumen were extracted using FIJI and analyzed with a custom Python script.

### 10. Statistical analysis

Tube diameter reproducibility was assessed by measuring internal (D□⍰ ⍰) and external (D□□⍰) diameters at three positions per tube (≥□1□mm apart), across N□≥□3 independently extruded tubes per condition. All quantitative results are expressed as mean□±□standard deviation (SD). Sample sizes and statistical tests are detailed in the respective figure legends.

Statistical analyses were performed using OriginPro (OriginLab, Northampton, MA; version 8) and Python 3.9 with standard libraries (*SciPy, statsmodels, pandas, matplotlib*). Normality of data distributions was assessed using the Shapiro–Wilk test. Depending on distribution and data structure, the following tests were applied:

- Two-group comparisons: unpaired or paired *t*-tests (parametric)
- Single-group comparisons: one-sample *t*-tests,
- Multiple group comparisons: one-way ANOVA with Tukey’s post-hoc test when parametric assumptions were met; otherwise, Kruskal–Wallis tests with Dunn’s correction.

For time-course analyses of proliferation markers (Ki-67 or PHH3), one-way ANOVA with Tukey’s test was used to compare all time points. Only non-significant comparisons (*p*□>□0.01) are annotated in figures to improve clarity. Complete test results are provided in Supplementary Tables S3, S4, and S5.

Significance thresholds were defined based on the actual distribution of p-values observed in this study. The following annotation scheme was used: **** p□<□0.0001, *** p□<□0.001, **p<0.01, *p<0.05, and ns = non-significant (p>0.01).

All protocols—including matrix preparation, extrusion parameters, culture conditions, and imaging settings—were applied uniformly across all endothelial cell types and time points. This methodological consistency enables robust, lineage-specific comparisons and ensures the reproducibility of key structural and functional readouts.

## Author contribution

Elsa Mazari-Arrighi: Conceptualization, Formal analysis, Investigation, Methodology, Visualization, Writing – original draft, Writing – review & editing. Adeline Boyreau: Investigation, Methodology. Laura Chaillot: Investigation, Methodology. Wilfried Souleyreau: Investigation, Methodology. Laetitia Andrique: Conceptualization, Investigation, Methodology. Thomas Mathivet: Conceptualization, Methodology, Supervision. Andreas Bikfalvi: Conceptualization, Formal analysis, Writing-editing, Methodology, Project administration, Resources, Supervision, Validation, Writing – review & editing. Pierre Nassoy: Conceptualization, Formal analysis, Funding acquisition, Methodology, Project administration, Resources, Supervision, Validation, Writing – review & editing.

## Data availability statement

The datasets supporting the findings of this study are available from the corresponding author upon reasonable request.

## Acknowledgments

We thank all members of the BiOf (Bioimaging and Optofluidics) team for their support. We are grateful to Naveen Mekhileri for synthesizing rhodamine-labeled gelatin, to Anirban Jana for assistance with the Incubascope, and Gaëlle Recher for her insightful scientific contributions. Multimodal microscopy was performed at the Bordeaux Imaging Center, a service unit of the CNRS-INSERM and Bordeaux University, member of the national infrastructure France BioImaging supported by the French National Research Agency (ANR-10-INBS-04). the Bordeaux Imaging Center (CNRS–INSERM, University of Bordeaux), member of the national infrastructure France BioImaging, for its continued support in multimodal microscopy.

