## SupplementaryFile for "A versatile microfluidic extrusion-based hydrogel platform for self-organization and long-term maintenance of engineered 3D lymphatic endothelium"

### Supplemental Figures

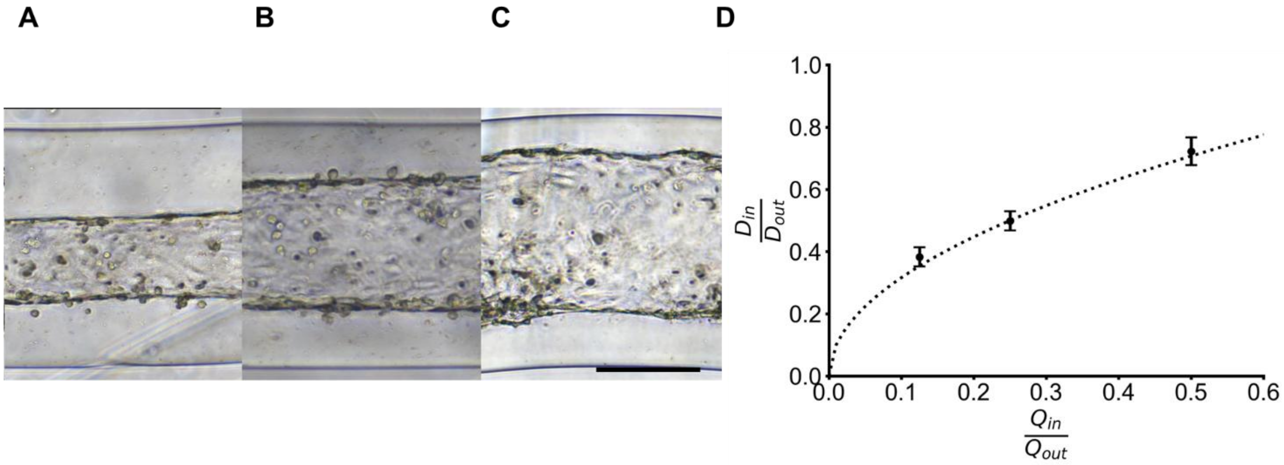


**Supplementary Figure S1**

**Fabrication of 3D lymphatic endothelium with tunable internal diameters using a 450-µm nozzle extrusion device.**

**(A–C)** Brightfield images of constructs produced with corresponding CS/IS/AL flow rate combinations of 0.25/0.25/3.5 mL/h **(A)**, 0.5/0.5/3 mL/h **(B)**, and 1/1/2 mL/h **(C)**. The internal diameter ($D_{in}$) expanded with the contribution of the HDLEC-laden core stream, reaching values above 300 µm. In contrast, the external diameter ($D_{out}$) remained stable (≈ 410 µm), as expected under the imposed constant total flow rate.

**(D)** Ratio $D_{in}/D_{out}$ as a function of the imposed flow rate ratio ($Q_{in}/Q_{out}$), with $Q_{in} = Q_{CS} + Q_{IS}$ and $Q_{out} = Q_{CS} + Q_{IS} + Q_{AL}$. The dotted curve corresponds to the theoretical prediction derived from volume conservation, following the square-root law:

$$\frac{D_{in}}{D_{out}} = \sqrt{\frac{Q_{in}}{Q_{out}}}$$

Black symbols represent experimental measurements of $D_{in}$ normalized to $D_{out}$, as also reported in **Fig. 1E**. Data are presented as mean ± standard deviation from ≥ 3 independent extrusion experiments per condition, each including ≥ 3 diameter measurements spaced by 1 mm to assess reproducibility.

**Scale bar:** 200 µm.

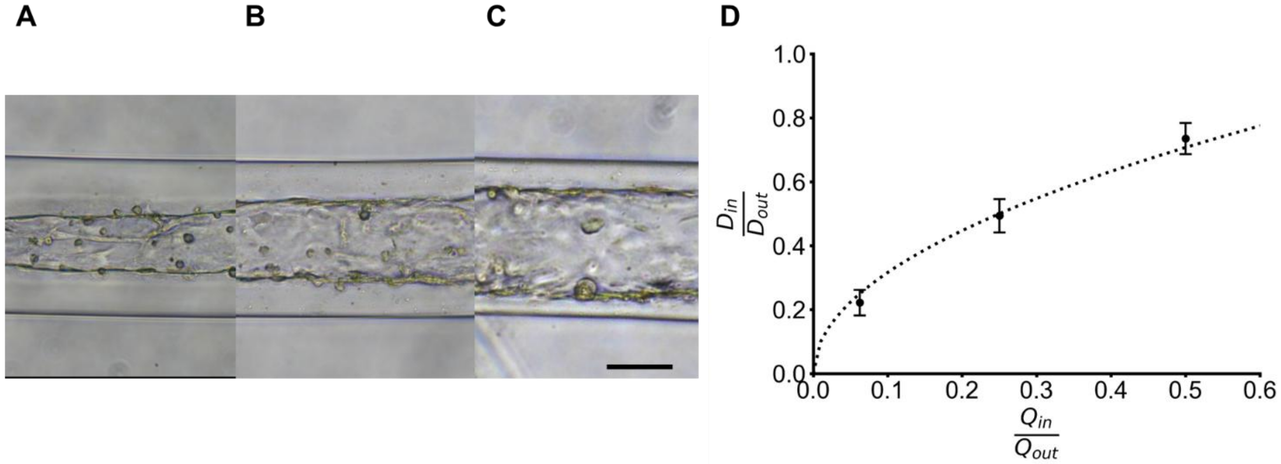


**Supplementary Figure S2**

**Fabrication of 3D lymphatic endothelium with tunable internal diameters using a 200-µm nozzle extrusion device.**

**(A–C)** Brightfield images of constructs produced with corresponding CS/IS/AL flow rate combinations of 0.15/0.15/3.7 mL/h **(A)**, 0.5/0.5/3 mL/h **(B)**, and 1/1/2 mL/h **(C)**. The internal diameter ($D_{in}$) expanded with the contribution of the HDLEC-laden core stream, ranging from ~50 to 150 µm. In contrast, the external diameter ($D_{out}$) remained stable (≈ 210 µm), as expected under the imposed constant total flow rate.

**(D)** Ratio $D_{in}/D_{out}$ as a function of the imposed flow rate ratio ($Q_{in}/Q_{out}$), with $Q_{in} = Q_{CS} + Q_{IS}$ and $Q_{out} = Q_{CS} + Q_{IS} + Q_{AL}$. The dotted curve corresponds to the theoretical prediction derived from volume conservation, following the square-root law:

$$\frac{D_{in}}{D_{out}} = \sqrt{\frac{Q_{in}}{Q_{out}}}$$

Black symbols represent experimental measurements of $D_{in}$ normalized to $D_{out}$, as also reported in **Fig. 1F**. Data are presented as mean ± standard deviation from ≥ 3 independent extrusion experiments per condition, each including ≥ 3 diameter measurements spaced by 1 mm to assess reproducibility.

**Scale bar:** 100 µm.

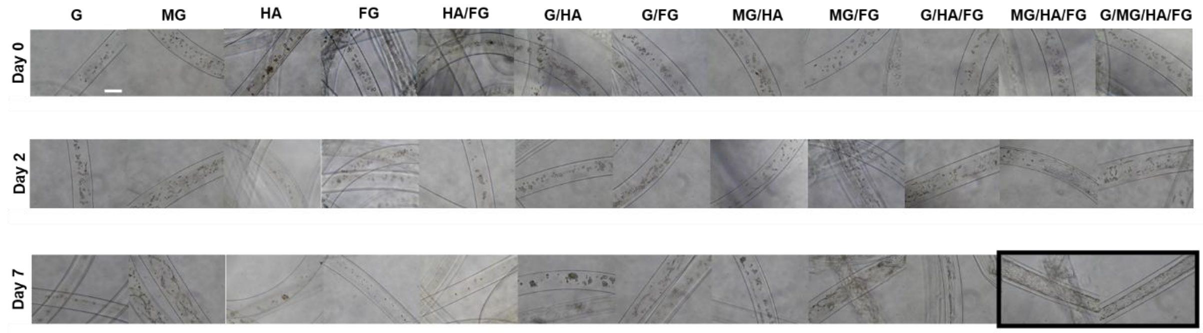


**Supplementary Figure S3**

**Matrix-driven HDLEC organization in tubular constructs.**

Brightfield and confocal images of HDLEC-laden artificial lymphatic endothelium formed in alginate-shelled tubes using various matrix compositions, cultured up to 7 days. Matrices were composed of 1-, 2-, 3-, or 4-component formulations based on gelatin (G), Matrigel (MG), hyaluronic acid (HA), and fibrinogen (FG). G and MG serve as scaffold-forming proteins [1–3], while HA and FG are bioactive factors known to influence lymphatic endothelial behavior [4–6].

(i) 1-component matrices: With G alone, HDLECs remained rounded and sparse over time, with minimal spreading between day 2 and day 7. With MG alone, cells rapidly migrated and extended along the alginate-core interface by day 2, consistent with Matrigel coating of alginate [2,3], forming aligned monolayers by day 7. HA alone failed to support any cell organization, lacking structural integrity. FG alone induced transient cord-like structures by day 2, which collapsed by day 7 due to insufficient mechanical support.

(ii) 2-component matrices: HA/FG yielded diffuse, disorganized cell patterns. G/HA and G/FG supported limited spreading without alignment. MG/HA permitted partial elongation, and MG/FG promoted early alignment that was not maintained by day 7.

(iii) 3-component matrices: MG/HA/FG induced dense, elongated cell assemblies reminiscent of endothelial cords, though complete monolayers were not consistently observed.

(iv) 4-component matrix (G/MG/HA/FG) enabled robust and reproducible formation of peripheral HDLEC monolayers by day 7

**Scale bar:** 200 μm.

**Supplementary Table S1**

**Morphological progression of lymphatic cell assemblies within tubular alginate constructs combining various natural ECM-derived protein components (gelatin, Matrigel, hyaluronic acid, and fibrinogen) at defined concentrations (w/v in total).** Lymphatic endothelial cells were encapsulated at $10^6$ cells/mL and monitored by brightfield microscopy at Day 2 and Day 7. While most protein combinations yield limited morphological definition over time, specific 3-component and 4-component formulations—MG/HA/FG and G/MG/HA/FG—support the emergence of hollow-like lymphatic self-organized tubular structures after 7 days of culture.

| Formulation | % Composition (in w/v) | Day | Observed Outcome |
|---|---|---|---|
| G | Gelatin 2% | Day 2 | Sparse distribution of individual cells |
| | | Day 7 | Low cell density; no visible tubular structures |
| MG | Matrigel 30% | Day 2 | Evenly distributed cells across the gel |
| | | Day 7 | Partial linear alignment; no continuous structures |
| HA | Hyaluronic acid 0.2% | Day 2 | Scattered, isolated cells |
| | | Day 7 | Diffuse signal; no visible organization |
| FG | Fibrinogen 0.2% | Day 2 | Dispersed cells; weak structural presence |
| | | Day 7 | Reduced signal intensity; no tubular features |
| HA/FG | HA 0.2%/FG 0.2% | Day 2 | Few grouped cells; unclear patterns |
| | | Day 7 | Isolated patches; no continuous structures |
| G/HA | G 2%/HA 0.2% | Day 2 | Local clustering and partial alignment |
| | | Day 7 | Short tubular segments visible |
| G/FG | G 2%/FG 0.2% | Day 2 | Thin, aligned lymphatic cell structures observed |
| | | Day 7 | Elongated lymphatic cell structures with visible continuity |
| MG/HA | MG 30%/HA 0.2% | Day 2 | Moderately dispersed cell pattern |
| | | Day 7 | Sparse cellular distribution; no visible organization |
| MG/FG | MG 30%/FG 0.2% | Day 2 | Initial arrangements of lymphatic cell structures |
| | | Day 7 | Outlined hollow-like tubular structures visible across the image |
| G/HA/FG | G 2%/HA 0.2%/FG 0.2% | Day 2 | Elongated cell groupings with limited extension |
| | | Day 7 | Extended lymphatic cell structures with partial branching |
| MG/HA/FG | MG 30%/HA 0.2%/FG 0.2% | Day 2 | Oriented cellular alignments observed |
| | | Day 7 | Outlined hollow-like tubular structures visible across the image |
| G/MG/HA/FG | G 2%/ MG 30%/HA 0.2%/FG 0.2% | Day 2 | Continuous aligned structures spanning across the gel |
| | | Day 7 | Outlined hollow-like tubular networks distributed throughout the gel |

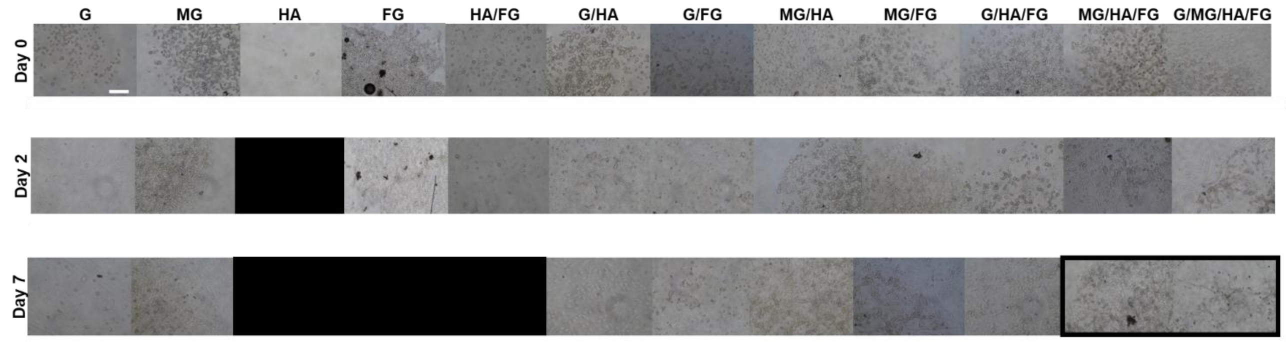


**Supplementary Figure S4**

**Matrix-driven HDLEC organization in droplet constructs.**

Brightfield images of HDLEC-laden droplets containing the same 1-, 2-, 3-, and 4-component formulations tested in Supplementary Figure S2-1, but polymerized without alginate to assess HDLEC behavior in unconfined environments.

In 1-component droplets, G and MG supported limited cell retention, with MG promoting early spreading and peripheral organization [2,3]. HA and FG (black image) alone failed to maintain structure, likely due to rapid diffusion and absence of scaffold formation.

Among 2-component formulations, MG/HA and MG/FG supported partial cord formation, while others remained poorly organized. The 3-component MG/HA/FG matrix led to compact multicellular assemblies by day 7. The 4-component G/MG/HA/FG matrix resulted in the most cohesive structures, forming organized, vessel-like cords even in the absence of confinement.

These complementary experiments emphasize the necessity of combining scaffold-forming [1–3] and bioactive components [4–6]. In unconfined environments, MG drives early spatial organization, while HA and FG enhance stability and morphogenesis. The 4-component formulation emerged as the most effective matrix to guide HDLECs into stable, self-organized vascular structures.

**Scale bar:** 200 µm.

**Supplementary Table S2**

**Morphological progression of lymphatic cell assemblies within droplet-based constructs combining natural ECM-derived protein components at defined concentrations (w/v in total).**

Lymphatic endothelial cells were encapsulated at $10^6$ cells/mL and monitored by brightfield microscopy at Day 2 and Day 7. Early culture stages revealed scattered or loosely grouped rounded cells, while several 3-component and 4-component formulations, notably MG/HA/FG and G/MG/HA/FG, promoted the formation of compact lymphatic cell structures.

| Formulation | % Composition (in w/v) | Day | Observed Outcome |
|---|---|---|---|
| G | Gelatin 2% | Day 2 | Moderate cell clustering; spherical cellular groupings observed |
| | | Day 7 | Low cell density; partial disaggregation of cell clusters |
| MG | Matrigel 30% | Day 2 | Centrally located cellular groupings; compact cell structures involved in lymphatic cord-like network assembly |
| | | Day 7 | Persistent compact cell structures; reduced peripheral cells |
| HA | Hyaluronic acid 0.2% | Day 2 | Image not available for this condition at this time point |
| | | Day 7 | Image not available for this condition at this time point |
| FG | Fibrinogen 0.2% | Day 2 | Scattered rounded cells; minimal cellular grouping |
| | | Day 7 | Image not available for this condition at this time point |
| HA/FG | HA 0.2%/FG 0.2% | Day 2 | Small cellular groupings; unevenly distributed |
| | | Day 7 | Image not available for this condition at this time point |
| G/HA | G 2%/HA 0.2% | Day 2 | Formation of isolated cell structures involved in lymphatic cord-like network assembly |
| | | Day 7 | Compact isolated cell structures with no observable continuity between them |
| G/FG | G 2%/FG 0.2% | Day 2 | Dense small aggregates of cells observed |
| | | Day 7 | Presence of compact round cell structures |
| MG/HA | MG 30%/HA 0.2% | Day 2 | Centrally located cellular groupings with homogeneous distribution |
| | | Day 7 | Increased proximity of cells; persistent multicellular groupings |
| MG/FG | MG 30%/FG 0.2% | Day 2 | Cell structures with central condensation observed |
| | | Day 7 | Well-defined, persistent cell structures involved in lymphatic cord-like network assembly |
| G/HA/FG | G 2%/HA 0.2%/FG 0.2% | Day 2 | Loose cellular groupings scattered across the matrix |
| | | Day 7 | More compact cell clusters; low structural regularity |
| MG/HA/FG | MG 30%/HA 0.2%/FG 0.2% | Day 2 | Prominent compact cell structures involved in lymphatic cord-like network assembly |
| | | Day 7 | Stable, well-defined cell structures involved in lymphatic cord-like network assembly |
| G/MG/HA/FG | G 2%/MG 30%/HA 0.2%/FG 0.2% | Day 2 | Numerous well-separated cell structures involved in lymphatic cord-like network assembly |
| | | Day 7 | Outlined, stable cell structures involved in lymphatic cord-like network assembly (highlighted in the figure) |

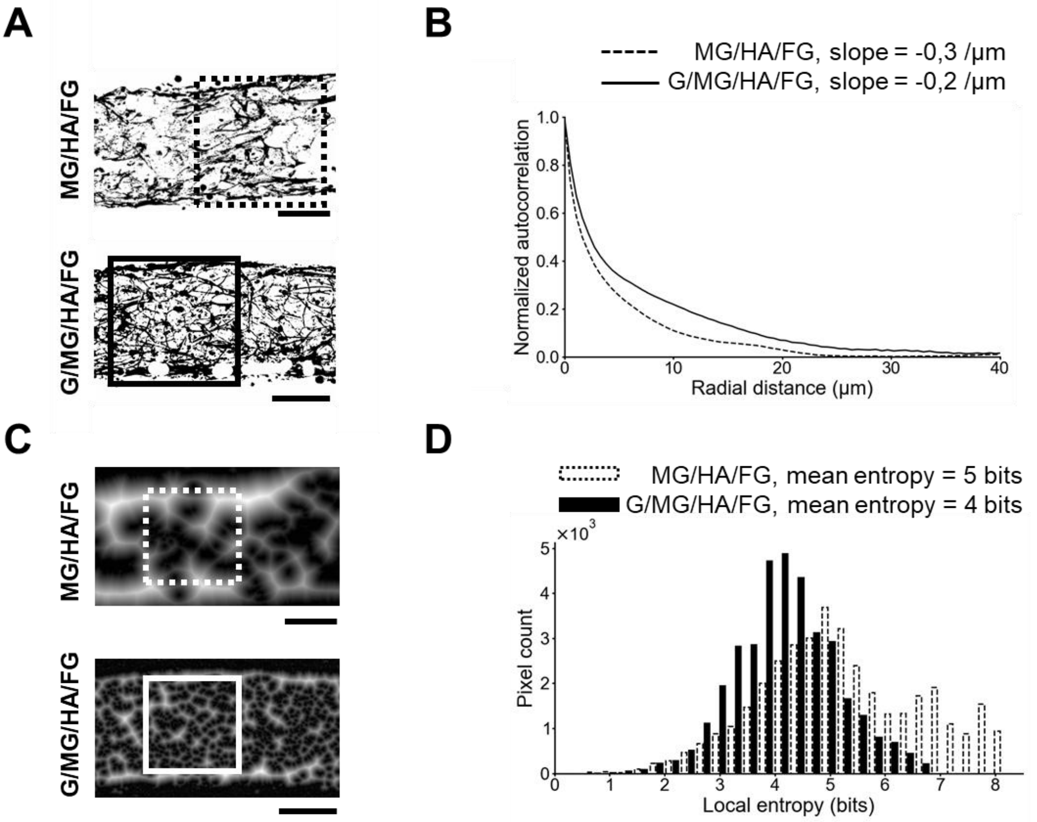


**Supplementary Figure S5**

**Characterization of actin cytoskeletal organization in MG/HA/FG and G/MG/HA/FG constructs at day 14.**

**(A)** Binarized images of actin filaments obtained after top-hat filtering and thresholding, illustrating the differences in filament continuity and density between MG/HA/FG (top) and G/MG/HA/FG (bottom) matrices.

**(B)** Radial autocorrelation profiles derived from the binarized images in panel **A**, showing a steeper decay for MG/HA/FG samples compared to G/MG/HA/FG.

**(C)** Nearest-neighbor distance histograms for actin structures, demonstrating greater spacing between filaments in MG/HA/FG constructs compared to G/MG/HA/FG.

**(D)** Local entropy measurements of actin organization within spatially defined regions of interest for MG/HA/FG and G/MG/HA/FG constructs, shown in panel **C** in blue and magenta respectively. These measurements further confirmed increased spatial disorder in MG/HA/FG compared to G/MG/HA/FG matrices (5 bits vs 4 bits).

**Scale bars:** 100 µm.

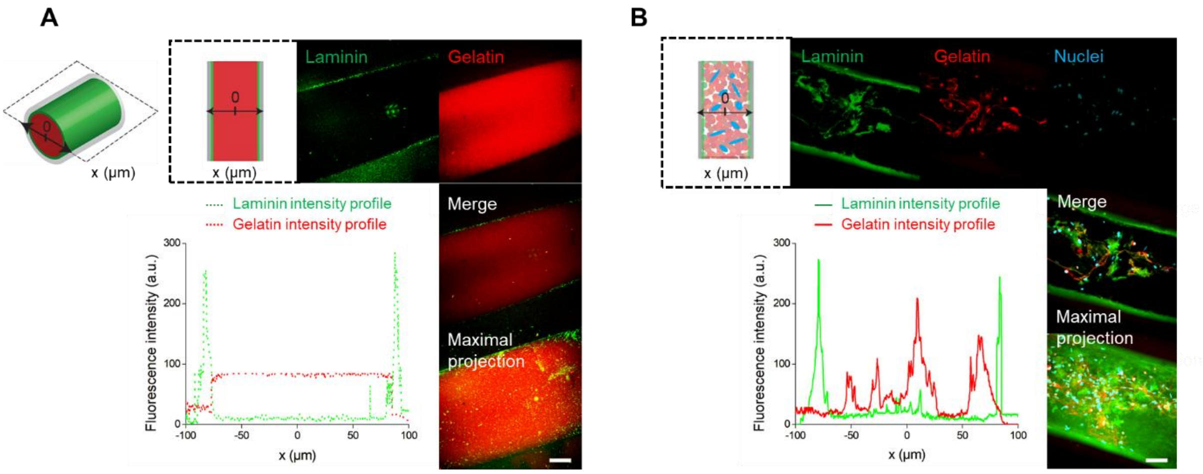


**Supplementary Figure S6**

**Matrix component distribution at day 1 with and without cells.**

**(A)** Cell-free constructs show uniform gelatin distribution (red) and peripheral laminin (green) coating the alginate wall, consistent with spontaneous Matrigel anchorage. [2,3].

**(B)** HDLEC-laden constructs display similar initial matrix organization, with no visible remodeling at this early time point.

**Scale bars:** 50 µm.

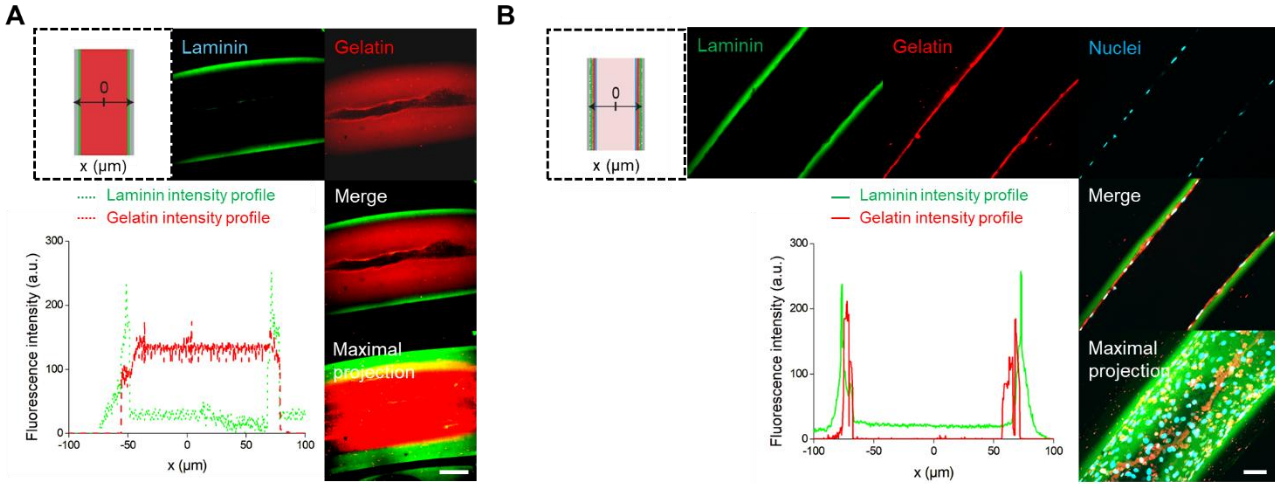


**Supplementary Figure S7**

**Matrix component distribution at day 14 with and without cells.**

**(A)** Cell-free constructs show uniform gelatin distribution (red) and peripheral laminin (green) coating the alginate wall, consistent with spontaneous Matrigel anchorage. [2,3].

**(B)** HDLEC-laden constructs display similar initial matrix organization, with no visible remodeling at this early time point.

**Scale bars:** 50 μm.

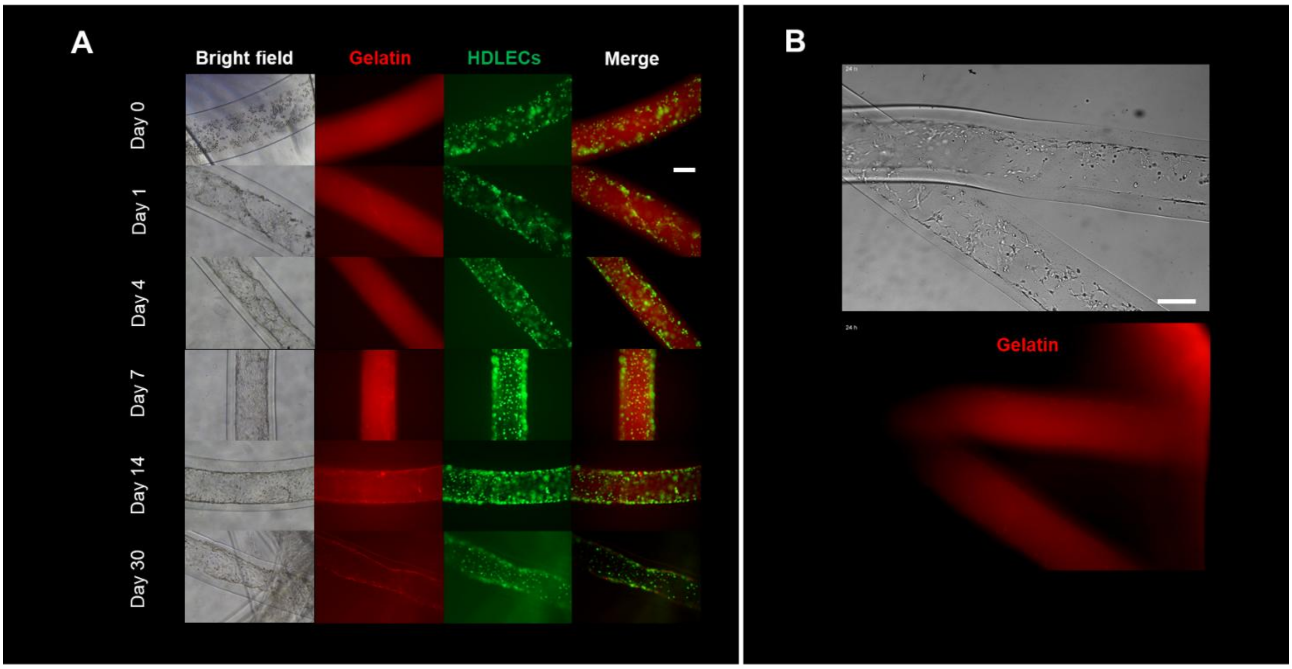


**Supplementary Figure S8**
**Time-course monitoring of gelatin remodeling and HDLEC organization within alginate-core constructs (day 0 to day 30).**
**(A)** Representative brightfield and fluorescence images of HDLEC-seeded constructs at days 0, 1, 4, 7, 14, and 30. Nuclei (green) correspond to HDLECs stably expressing H2B-GFP. The hydrogel core contains rhodamine-labeled gelatin ($G_{rh}$; red) as part of the $G_{rh}$/MG/HA/FG matrix.
**(B)** Fluorescence and structural changes over time reflect progressive cell organization and gelatin remodeling.
**Scale bar:** 100 μm.

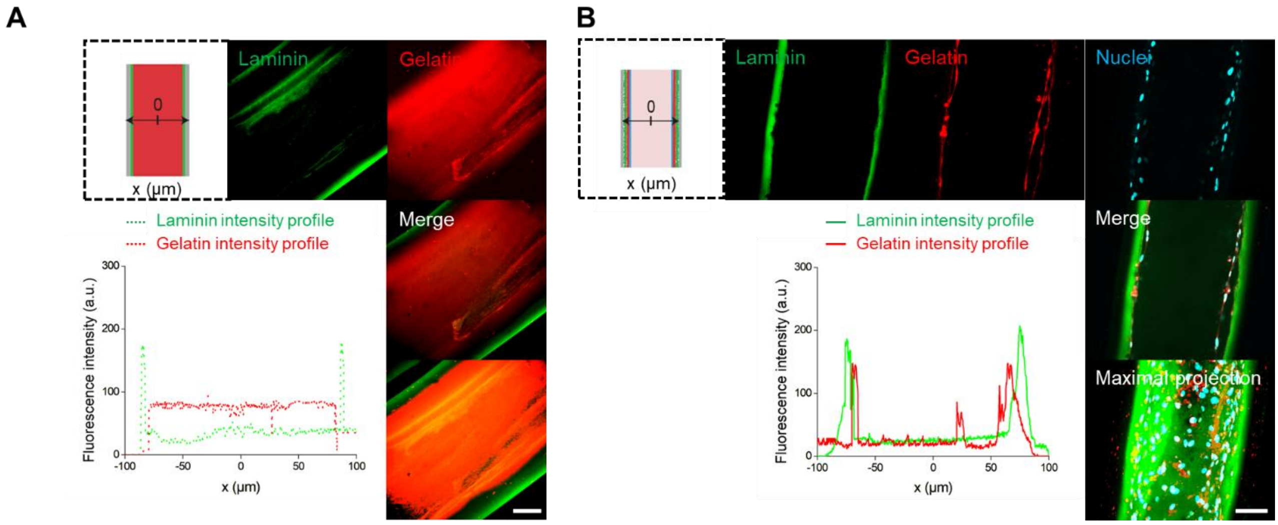


**Supplementary Figure S9**

**Matrix component distribution at day 30 with and without cells.**

**(A)** Cell-free constructs show uniform gelatin distribution (red) and peripheral laminin (green) coating the alginate wall, consistent with spontaneous Matrigel anchorage [2,3].

**(B)** HDLEC-laden constructs display similar initial matrix organization, with no visible remodeling at this early time point.

**Scale bars:** 50 µm.

| Comparison | p-value | Significance |
| --- | --- | --- |
| Day1 vs Day14 | <1e-16 | **** |
| Day1 vs Day21 | <1e-16 | **** |
| Day1 vs Day3 | <1e-16 | **** |
| Day1 vs Day30 | <1e-16 | **** |
| Day1 vs Day5 | <1e-16 | **** |
| Day1 vs Day7 | 0.0004 | *** |
| Day14 vs Day21 | 0.8258 | ns |
| Day14 vs Day3 | <1e-16 | **** |
| Day14 vs Day30 | 0.1148 | ns |
| Day14 vs Day5 | <1e-16 | **** |
| Day14 vs Day7 | 0.9113 | ns |
| Day21 vs Day3 | <1e-16 | **** |
| Day21 vs Day30 | 0.7812 | ns |
| Day21 vs Day5 | <1e-16 | **** |
| Day21 vs Day7 | 0.1770 | ns |
| Day3 vs Day30 | <1e-16 | **** |
| Day3 vs Day5 | <1e-16 | **** |
| Day3 vs Day7 | <1e-16 | **** |
| Day30 vs Day5 | <1e-16 | **** |
| Day30 vs Day7 | 0.0062 | ** |
| Day5 vs Day7 | <1e-16 | **** |

**Supplementary Table S3**
**Statistical analysis of PHH3-positive cell percentages across the culture period (Days 1 to 30), corresponding to Fig. 3B.**
Mitotic activity was assessed by quantifying the percentage of PHH3-positive nuclei at each culture time point. One-way ANOVA followed by Tukey's post hoc multiple comparisons test was used to identify statistically significant differences between all pairs of time points. The table provides the adjusted p-values along with corresponding levels of significance (**** p < 0.0001; *** p < 0.001; ** p < 0.01; *p<0,05; ns = not significant).
To optimize readability of the graph (**Fig. 3B**), only comparisons with p > 0.05 ("ns") are explicitly shown. All significant comparisons are listed in this table for comprehensive reference.

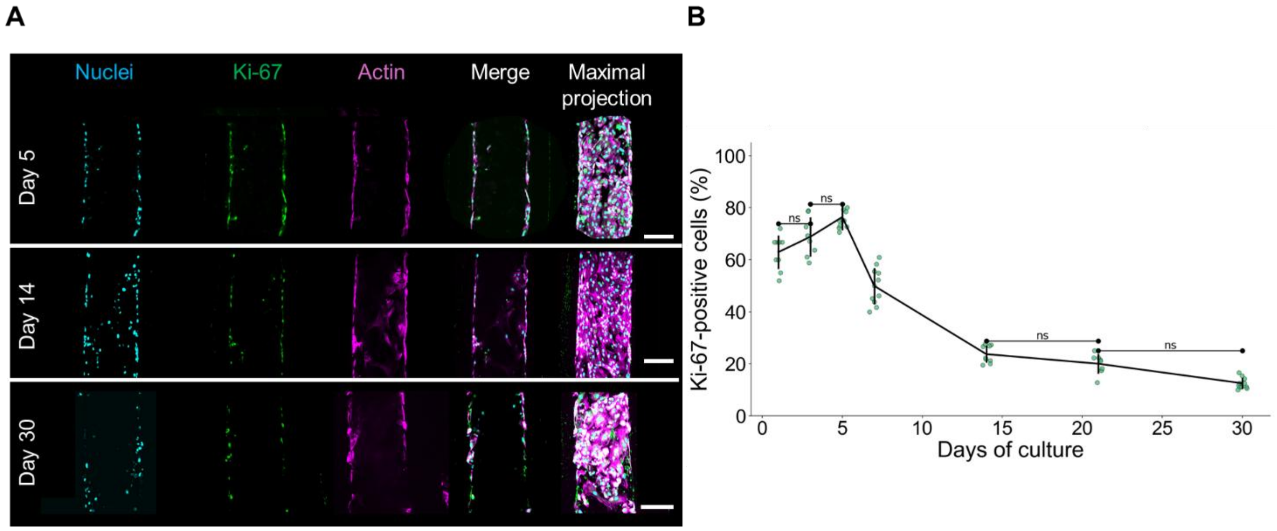


**Supplementary Figure S10**

**Time-dependent Ki-67 expression in 3D endothelial constructs.**

**(A)** Confocal equatorial images showing immunostaining for nuclei (cyan), Ki-67 (green), and actin (magenta) at day 5, day 14, and day 30. Merged equatorial as well as maximal projection views highlight spatial localization of Ki-67-positive cells and changes in signal density.

**(B)** Scatter plot displaying individual data points representing the percentage of Ki-67-positive cells relative to total nuclei over time. Each dot corresponds to an individual measurement; vertical bars indicate the mean ± standard deviation for each time point, N ≥ 3 constructs per time point at least. Only pairwise comparisons with $p > 0.05$ are explicitly shown on the graph (annotated with ns, *, etc.). All other comparisons are shown in **Table S4**.

**Scale bars:** 100 µm.

| Comparison | p-value | Significance |
|---|---|---|
| 1 vs 3 | 0.2777 | ns |
| 1 vs 5 | <1e-16 | **** |
| 1 vs 7 | <1e-16 | **** |
| 1 vs 14 | <1e-16 | **** |
| 1 vs 21 | <1e-16 | **** |
| 1 vs 30 | <1e-16 | **** |
| 3 vs 5 | 0.0489 | ns |
| 3 vs 7 | <1e-16 | **** |
| 3 vs 14 | <1e-16 | **** |
| 3 vs 21 | <1e-16 | **** |
| 3 vs 30 | <1e-16 | **** |
| 5 vs 7 | <1e-16 | **** |
| 5 vs 14 | <1e-16 | **** |
| 5 vs 21 | <1e-16 | **** |
| 5 vs 30 | <1e-16 | **** |
| 7 vs 14 | <1e-16 | **** |
| 7 vs 21 | <1e-16 | **** |
| 7 vs 30 | <1e-16 | **** |
| 14 vs 21 | 0.8021 | ns |
| 14 vs 30 | 0.004 | ** |
| 21 vs 30 | 0.0507 | ns |

**Supplementary Table S4**
**Statistical summary of Ki-67-positive cell percentages across the culture timeline (Days 1 to 30), corresponding to Fig. S10B.**
To analyze the temporal dynamics of proliferating cells, the percentage of Ki-67-positive nuclei was quantified at each time point. Statistical significance was evaluated using one-way ANOVA followed by Tukey's post-hoc test, allowing pairwise comparisons between all days. The table lists the adjusted p-values along with their respective significance indicators (**** p < 0.0001; *** p < 0.001; ** p < 0.01; *p<0,05; ns = not significant).

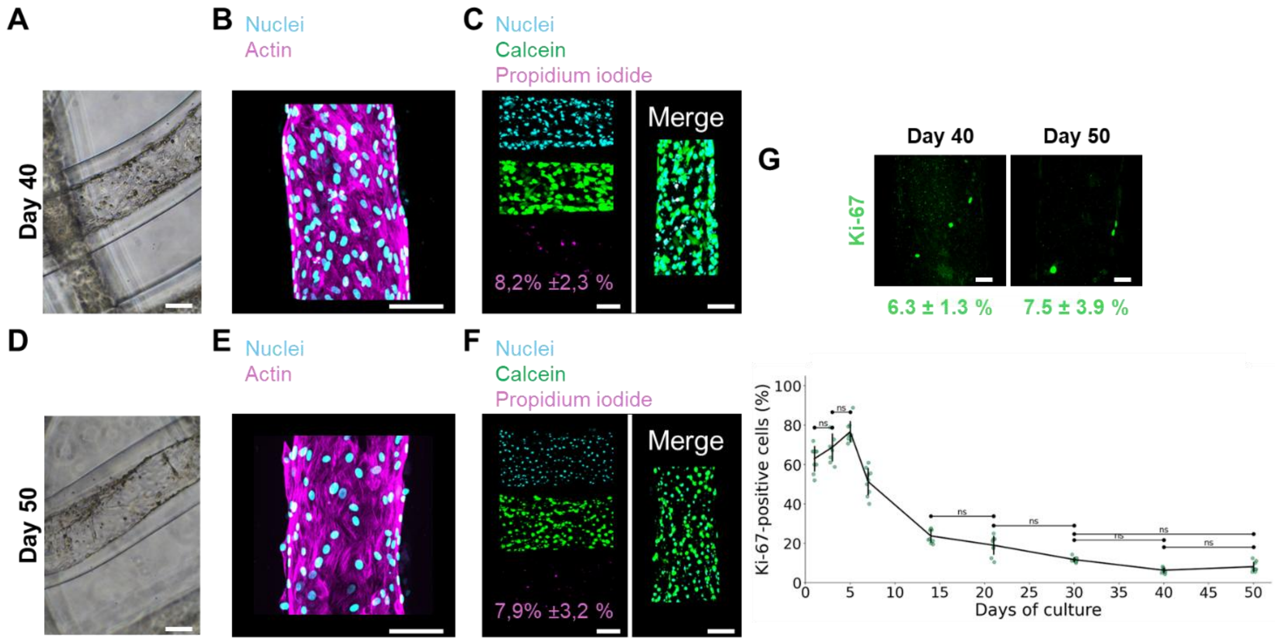


**Supplementary Figure S11**

**Extended analysis of structural stability, viability, and proliferative activity in artificial 3D lymphatic endothelium at days 40 and 50 under static culture conditions.**

**(A, B)** Brightfield images of constructs at days 40 **(A)** and 50 **(B)**, showing maintained tubular morphology and cell density.

**(D, E)** Confocal maximum intensity projections stained for nuclei (cyan) and actin (magenta) at days 40 **(D)** and 50 **(E)**, revealing persistent cortical actin organization and regular nuclear distribution.

**(C, F)** Viability assays at days 40 **(C)** and 50 **(F)**, with nuclei (cyan), live cells (green), and dead cells (magenta) staining. Quantification indicates low and stable percentages of dead cells (~8%) at both time points.

**(G)** Ki-67 immunostaining at days 40 and 50 showing minimal proliferative activity. Dot plot quantifies individual values of Ki-67 positive cells. N ≥ 3 per tube for viability and Ki-67 analysis. Each dot corresponds to an individual measurement; vertical bars indicate the mean ± standard deviation for each time point. Only pairwise comparisons with $p > 0.05$ are explicitly shown on the graph (annotated with ns). All other comparisons are shown in **Table S5**.

**Scale bars:** 100 µm **(A–F)**, 50 µm **(G).**

| Comparison | p-value | Significance |
|---|---|---|
| Day1 vs Day14 | <1e-16 | **** |
| Day1 vs Day21 | <1e-16 | **** |
| Day1 vs Day3 | 0.2267 | ns |
| Day1 vs Day30 | <1e-16 | **** |
| Day1 vs Day40 | <1e-16 | **** |
| Day1 vs Day5 | <1e-16 | **** |
| Day1 vs Day50 | <1e-16 | **** |
| Day1 vs Day7 | 1.000e-04 | *** |
| Day14 vs Day21 | 0.5133 | ns |
| Day14 vs Day3 | <1e-16 | **** |
| Day14 vs Day30 | 0.004 | *** |
| Day14 vs Day40 | <1e-16 | **** |
| Day14 vs Day5 | <1e-16 | **** |
| Day14 vs Day50 | <1e-16 | **** |
| Day14 vs Day7 | <1e-16 | **** |
| Day21 vs Day3 | <1e-16 | **** |
| Day21 vs Day30 | 0.0543 | ns |
| Day21 vs Day40 | <1e-16 | **** |
| Day21 vs Day5 | <1e-16 | **** |
| Day21 vs Day50 | 0.0004 | *** |
| Day21 vs Day7 | <1e-16 | **** |
| Day3 vs Day30 | <1e-16 | **** |
| Day3 vs Day40 | <1e-16 | **** |
| Day3 vs Day5 | 0.0589 | ns |
| Day3 vs Day50 | <1e-16 | **** |
| Day3 vs Day7 | <1e-16 | **** |
| Day30 vs Day40 | 0.3295 | ns |
| Day30 vs Day5 | <1e-16 | **** |
| Day30 vs Day50 | 0.8225 | ns |
| Day30 vs Day7 | <1e-16 | **** |
| Day40 vs Day5 | <1e-16 | **** |
| Day40 vs Day50 | 0.9969 | ns |
| Day40 vs Day7 | <1e-16 | **** |
| Day5 vs Day50 | <1e-16 | **** |
| Day5 vs Day7 | <1e-16 | **** |
| Day50 vs Day7 | <1e-16 | **** |

**Supplementary Table S5**
**Statistical analysis of Ki-67-positive cell percentages across culture time from Day 1 to Day 50, related to Fig. S11G.**

Cell proliferation dynamics were quantified by assessing the percentage of Ki-67-positive nuclei at each time point. One-way ANOVA followed by Tukey’s multiple comparisons test was used to evaluate statistical differences between all pairs of time points. The table reports adjusted p-values and corresponding significance levels (****$p < 0.0001$; *** $p < 0.001$; ** $p < 0.01$; *p<0,05; ns = not significant).

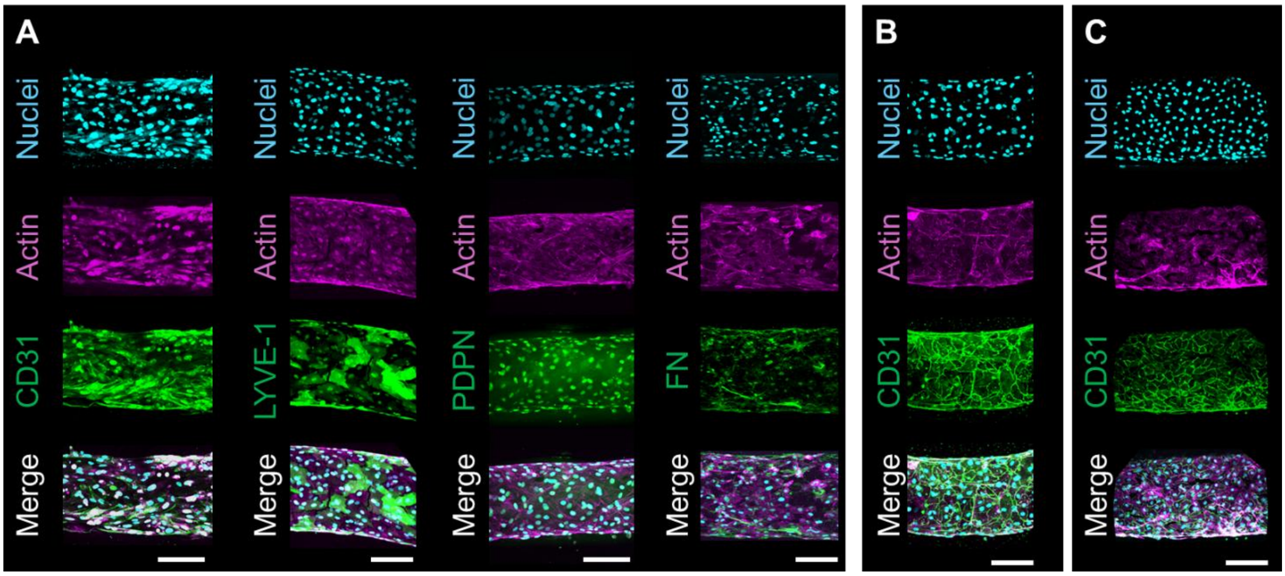


**Supplementary Figure S12**

**Day-14 lineage-specific marker expression in 3D endothelial constructs.**

**(A–C)** Confocal imaging of HDLEC- (**A**), HDMEC- **(B)**, and HUVEC-based (**C)** constructs at day 14. HDLECs exhibit expression of CD31, LYVE-1, PDPN, and fibronectin (FN), while HDMECs and HUVECs express only CD31. These observations show that lineage-specific identity and ECM remodeling are established during the first two weeks of culture.

**Scale bars:** 100 µm.

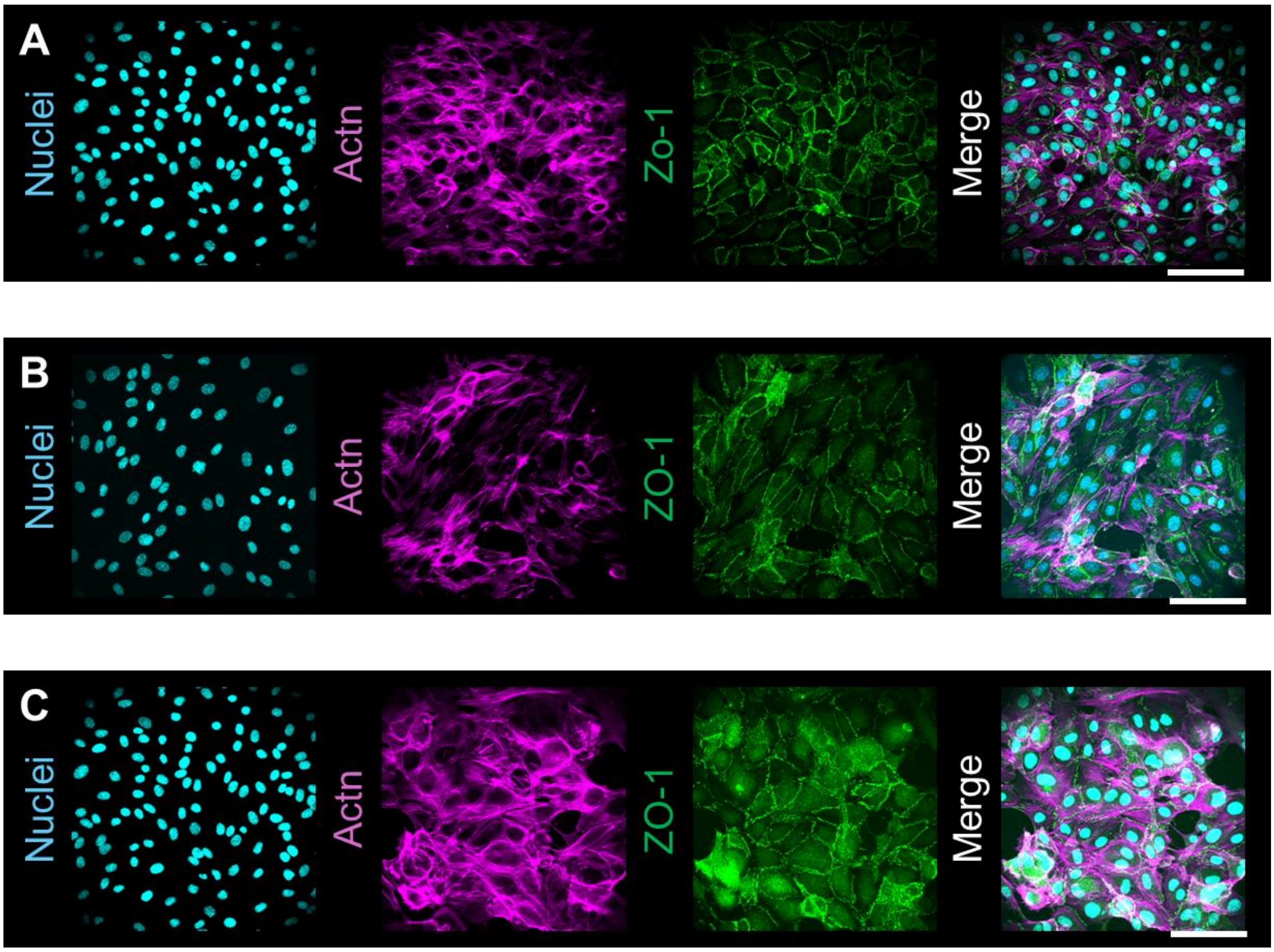


**Supplementary Figure S13**
**ZO-1 expression in 2D cultures of endothelial cells.**
**(A–C)** HDLECs **(A)**, HDMECs **(B),** and HUVECs **(C)** cultured on 2D substrates all express ZO-1 at intercellular junctions under identical conditions.
**Scale bars:** 100 µm.

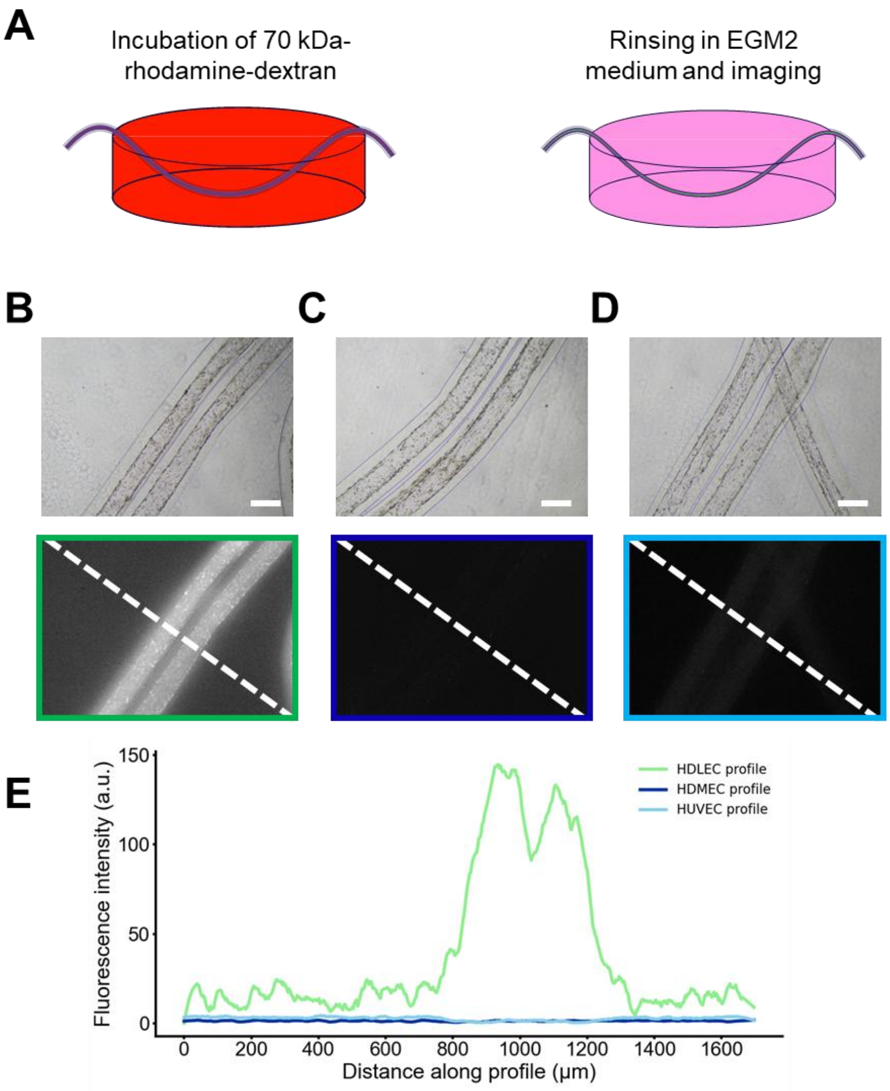


**Supplementary Figure S14**

**Schematic and fluorescence imaging of the 70 kDa dextran permeability assay.** **(A)** Schematic representation of the permeability assay setup. Tubes were incubated for 30 minutes in a rhodamine-labeled 70 kDa dextran solution, with both extremities carefully positioned outside the culture well to ensure selective exposure of the lateral surface to the tracer. This configuration restricted dextran access to the basal side of the endothelial layer. After incubation, samples were thoroughly rinsed and the dextran solution was replaced with fresh culture medium to stop further diffusion and clear residual tracer from the exterior. Fluorescence imaging was then performed to assess intraluminal accumulation.

**(B–D)** Low-magnification fluorescence images of HDLEC- **(B)**, HDMEC- **(C)**, and HUVEC-based **(D)** endothelial monolayers at day 30. Only HDLEC-derived endothelium showed luminal accumulation of the tracer. In contrast, HDMEC- and HUVEC-derived endothelia excluded the tracer.

**(E)** Fluorescence intensity profiles measured across the lumens shown in **(B–D)**, illustrating differential tracer penetration across the three endothelial types. **Scale bars:** 200 μm.

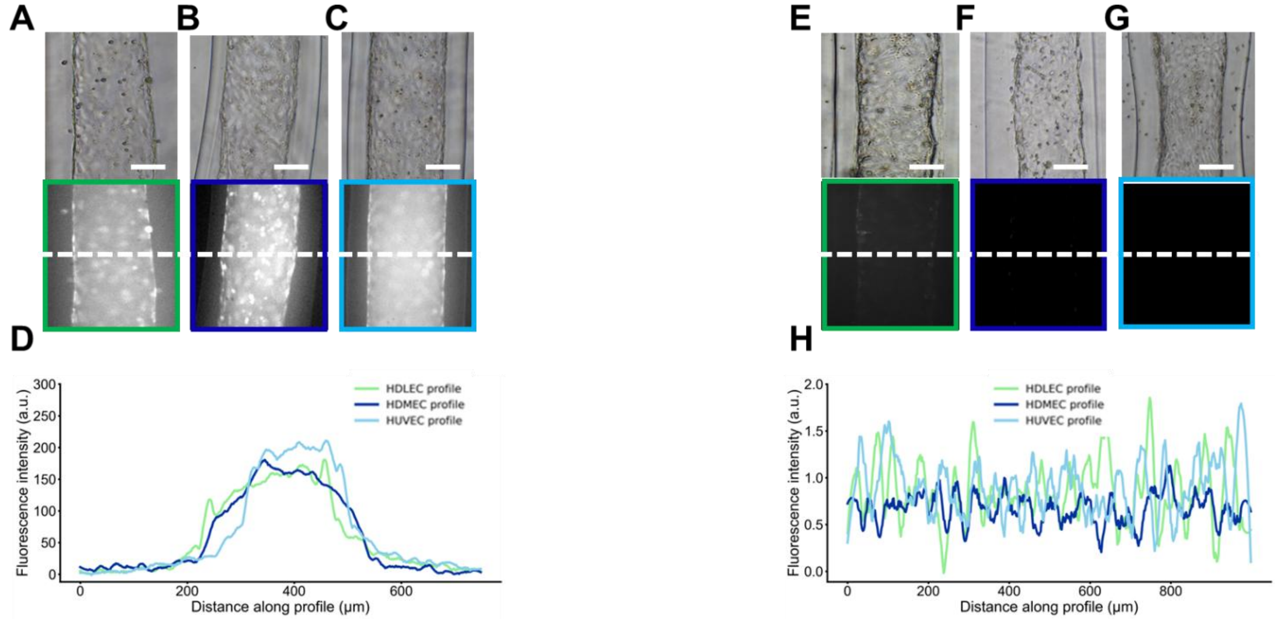


**Supplementary Figure S15**

**Size-dependent permeability of endothelial constructs.**

**(A–D)** Permeability assay using 3 kDa FITC-dextran. Constructs composed of HDLECs **(A)**, HDMECs **(B)**, and HUVECs **(C)** were incubated with 3 kDa fluorescent tracer. Fluorescence imaging revealed tracer accumulation in the lumen of all constructs, indicating permeation across the alginate shell and endothelial layers. Panel D shows intensity profiles across tube cross-sections, confirming comparable tracer penetration in all conditions.

**(E–H)** Permeability assay using 500 kDa rhodamine-dextran. Constructs composed of HDLECs **(E)**, HDMECs **(F)**, and HUVECs **(G)** were incubated with high-molecular-weight tracer. No luminal fluorescence signal was detected in any condition, as shown in intensity profiles **(H)**, confirming that 500 kDa dextran is effectively excluded by both the surrounding alginate layer and the endothelial barrier [7].

**Scale bars:** 100 µm.

### Supplemental movies

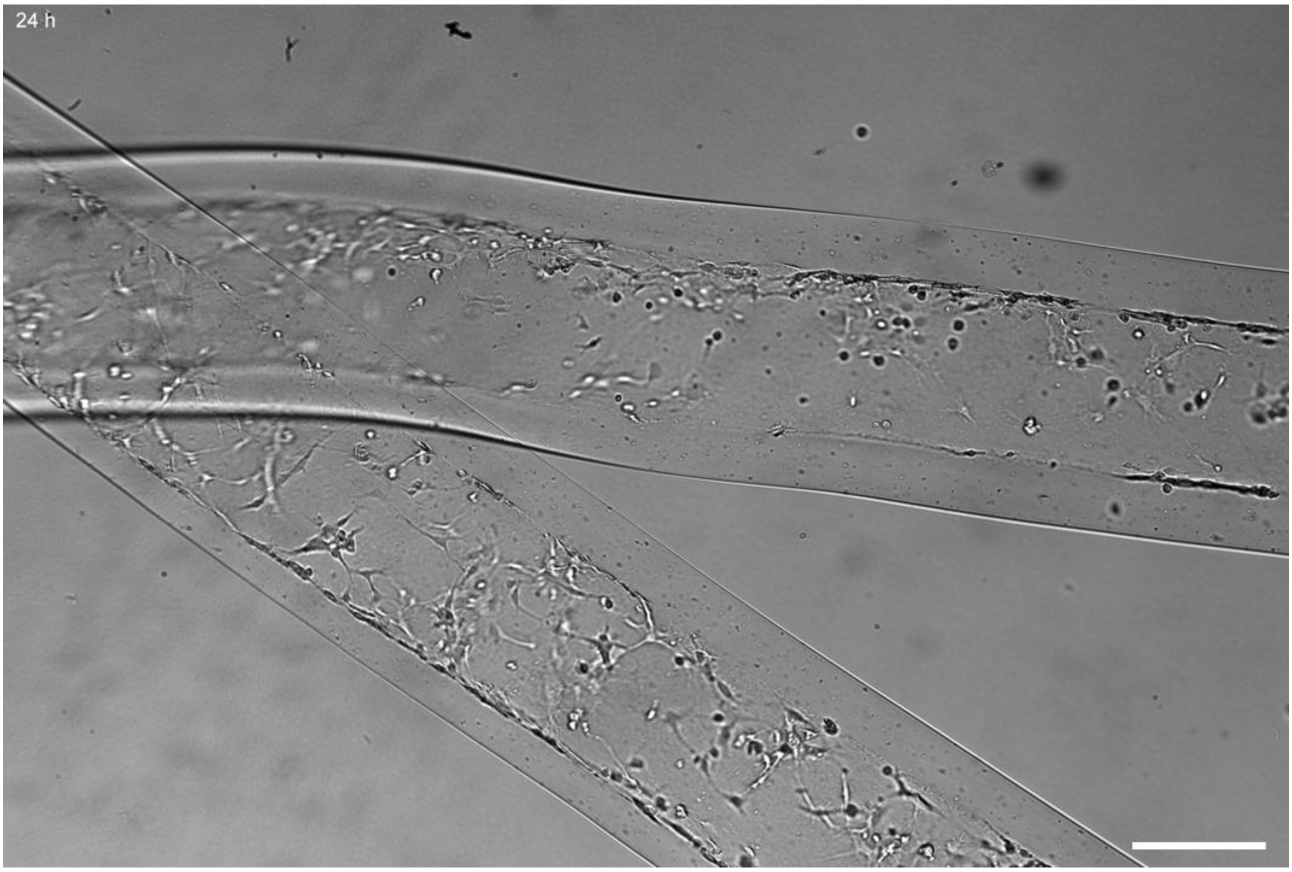


**Supplementary Movie S1**
**Time-lapse monitoring of HDLEC organization and gelatin remodeling in four-component matrices ($G_{rh}$/MG/HA/FG) containing rhodamine-labeled gelatin ($G_{rh}$).** Brightfield time-lapse sequence showing the dynamic behavior of HDLECs embedded within the hydrogel core. One image was acquired every 3 hours using an Incubascope system (see [8]). Progressive lymphatic endothelial reorganization is observed throughout a 28-day culture period.
**Scale bar:** 100 µm.

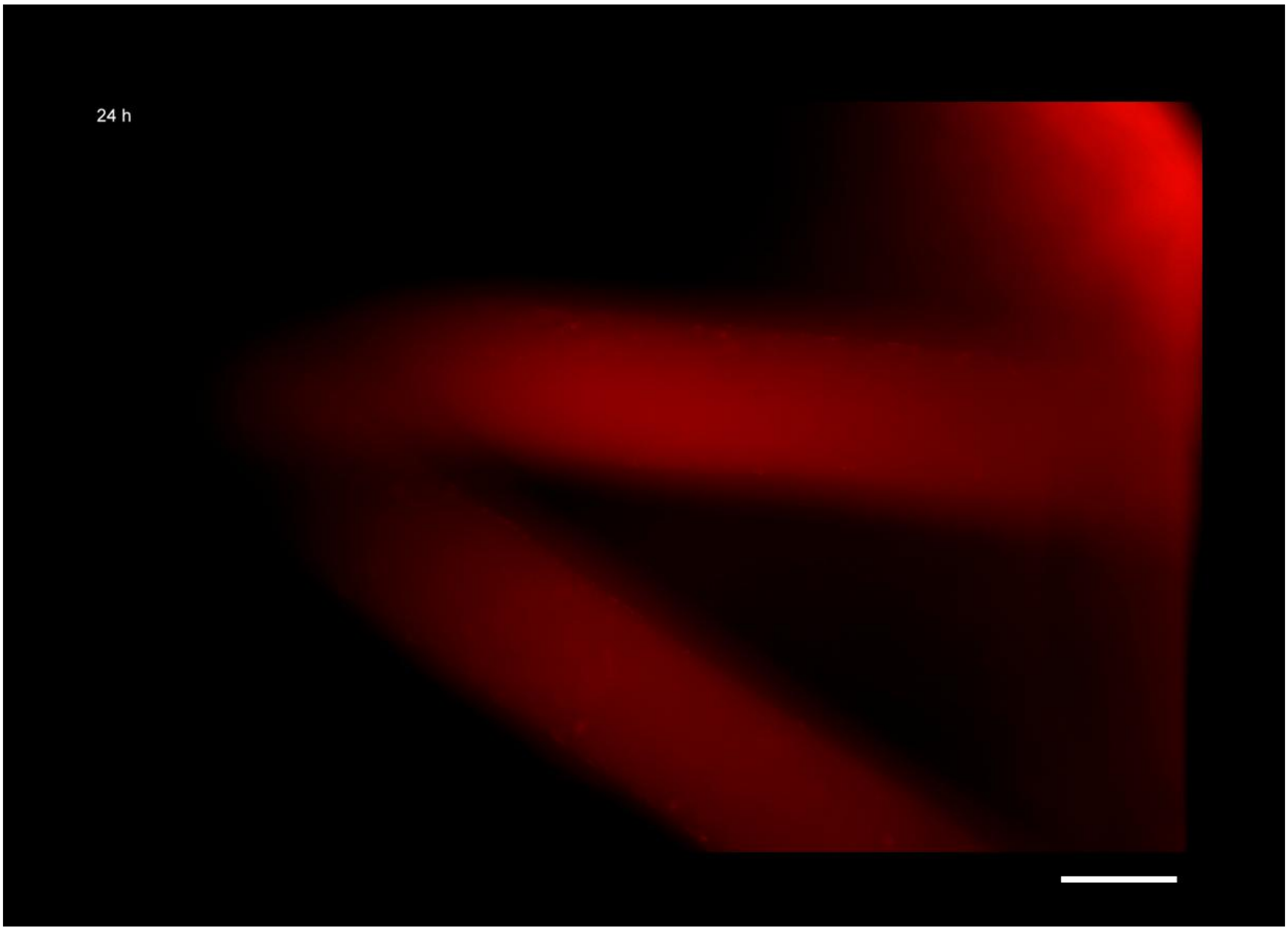


**Supplementary Movie S2**
**Time-lapse monitoring of gelatin remodeling in four-component matrices seeded with HDLECs.**
Live fluorescence imaging from day 1 to day 29 showing rhodamine-labeled gelatin ($G_{rh}$, red) within a $G_{rh}$/MG/HA/FG matrix seeded with HDLECs. One image was acquired every 3 hours using an Incubascope system (see [8]). Progressive loss of $G_{rh}$ signal becomes apparent around day 13, consistent with gelatin degradation and matrix remodeling by embedded cells.
**Scale bar:** 100 µm.
